# Fine tuning of Rpb4 phosphorylation modulates Rpb4/7 stoichiometry within yeast RNA polymerase II and regulates gene expression

**DOI:** 10.1101/2023.12.15.571801

**Authors:** Araceli González-Jiménez, María del Carmen González-Jiménez, Antonio Jordán-Pla, Ithaisa Medina, Manuel J. Alfonso, Stephen Richard, Carlos Fernández-Tornero, Mordechai Choder, Olga Calvo

## Abstract

RNA polymerase II (RNAPII) consists of 12 subunits, including Rpb4 and Rpb7, which form a heterodimeric stalk domain. In yeast, assembly of Rpb4/7 with the 10-subunit “core” is substoichiometric. Aside from its role in transcription, Rpb4/7 also binds to mRNAs co-transcriptionally (mRNA imprinting), exits the nucleus together with the mRNA, and modulates subsequent cytoplasmic processes like translation and degradation. However, the mechanisms underlying Rpb4/7 functions and its association with RNAPII remain poorly understood. In this study, we identify five phosphorylation sites in Rpb4 that regulate various steps of mRNA and snoRNA biogenesis, including transcription elongation and the polyadenylation step. We also provide evidence that the kinase Hrr25 and the phosphatase Fcp1, known to target the carboxyl-terminal domain (CTD) of Rpb1 and participate in processes at gene 3′-ends, coordinate Rpb4 phosphorylation. Inefficient Rpb4 phosphorylation leads to increased Rpb4 association with RNAPII, particularly at gene 3′-ends, suggesting that Rpb4-P promotes stalk dissociation from RNAPII either before or during transcription termination. Collectively, our findings underscore the pivotal role of dynamic Rpb4 phosphorylation in regulating Rpb4/7 stoichiometry as well as transcription elongation and termination, with an impact on mRNA imprinting.

## INTRODUCTION

RNAPII consists of 12 subunits, from Rpb1 to Rpb12, with Rpb12 being the smallest one. Rpb1 contains a flexible and unstructured carboxyl-terminal domain (CTD). It is a unique evolutionarily conserved domain consisting of a tandem repetition of the Tyr1-Ser2-Pro3-Thr4-Ser5-Pro6-Ser7 (YSPTSPS) sequence (Zhang and Corden 1991; Chapman et al. 2008). The CTD is subject to numerous post-translational modifications (PTM). It can be phosphorylated (Zhang and Corden 1991; Schuller et al. 2016; Suh et al. 2016), methylated (Sims et al. 2011), acetylated (Schroder et al. 2013), and glycosylated (Ranuncolo et al. 2012), with phosphorylation being the main PTM. Besides, isomerization of the two prolines residues regulates CTD phosphorylation (Hanes 2014).

CTD phosphorylation is widely recognized as essential for regulating co-transcriptional pre-mRNA processing and overall gene expression. Although five of the seven CTD residues can be phosphorylated (Tyr1, Ser2, Thr4, Ser5, and Ser7) (Buratowski 2003; Hsin and Manley 2012; Yurko and Manley 2018), it is mostly phosphorylation of Ser2 and Ser5 that contribute to the output of RNAPII phosphorylation levels (Schuller et al. 2016; Suh et al. 2016). Phosphorylation/dephosphorylation of the CTD is dynamically regulated during the transcription cycle following an extraordinarily complex code, named the “CTD code” (Buratowski 2003). This code promotes and coordinates the timely recruitment of many proteins that recognize specific and distinct CTD phosphorylation patterns. In this manner, the CTD acts as a scaffolding platform able to assemble or release factors involved in many aspects of mRNA biogenesis, mediating its connection to other nuclear processes (Perales and Bentley 2009; Calvo and García 2012; Hsin and Manley 2012; Harlen and Churchman 2017). After the discovery of Rpb1-CTD and its role in pre-mRNA processing, its PTM, and specifically phosphorylation, have been extensively studied. This is not the case for other RNAPII subunits, although a number of proteomic studies have identified residues present in subunits other than Rpb1 that are susceptible to PTM (Richard et al. 2021). We recently compiled all RNAPII phospho-sites identified in proteomic studies and found that, indeed, all human RNAPII subunits contain phospho-sites, while in *Saccharomyces cerevisiae* there are no phospho-sites in Rpb7 and Rpb11 (Gonzalez-Jimenez et al. 2021). However, whether phosphorylation of RNAPII subunits other than Rpb1 has any biological role remains still unknown.

We centered our attention on Rpb4 that, together with Rpb7, forms the stalk module, which is conserved from yeast to humans and has orthologous modules in RNAPI, RNAPIII, and archaeal RNAP (Grohmann and Werner 2011; Werner and Grohmann 2011). Rpb4/7 protrudes from the core complex near the CTD and the RNA exit channel (Fig. 1B) (Armache et al. 2005), a position with potential interaction with CTD-modifying enzymes, nascent RNAs, pre-mRNA processing factors, and, possibly, other regulatory factors (Allepuz-Fuster et al. 2014; Garavis et al. 2017; Allepuz-Fuster et al. 2019; Calvo 2020). In higher eukaryotes, Rpb4 and Rpb7 are essential subunits (Zhao et al. 2012; Sharma and Kumari 2013). In *S. cerevisiae*, although Rpb7 is essential, deletion of *RPB4* only results in cell lethality under certain conditions (Choder and Young 1993). Rpb4/7 associates co-transcriptionally with nascent mRNAs in the nucleus and accompanies them to the cytoplasm in a process termed mRNA imprinting (Choder 2011; Dahan and Choder 2013; Duek et al. 2018; Chalabi Hagkarim and Grand 2020). In the cytoplasm, Rpb4/7 stimulates mRNA translation initiation (Duek et al. 2018) and subsequent decay (Lotan et al. 2005; Lotan et al. 2007; Goler-Baron et al. 2008). After RNA decay, Rpb4/7 returns to the nucleus, completing a cyclical process (Haimovich et al. 2013). Rpb4/7 regulates all major processes that mRNAs undergo via mRNA imprinting. How Rpb4/7 participates in spatially and temporally separated processes regulating gene expression is explained in part by its ability to interact with different nuclear and cytosolic complexes, but how these interactions are regulated is a fundamental question that remains unresolved. Recently, it has been shown that numerous PTM of Rpb4/7, in particular Rpb4 methylation, acetylation, and ubiquitylation, link transcription to post-transcriptional processes and are important for maintaining proper mRNA level—a phenomenon known as “mRNA buffering” (Richard et al. 2021). It was proposed that the numerous PTM combinations serve as a language for communication across stages of the gene expression system (Richard et al. 2021). Here, we present data demonstrating that Rpb4 is a phospho-protein and that phosphorylated Rpb4 (Rpb4-P) levels regulate gene transcription and Rpb4 stoichiometry within RNAPII, preferentially at the 3′-ends of genes. Moreover, we have identified Hrr25 kinase and Fcp1 phosphatase as regulators of Rpb4-P levels. Interestingly, Hrr25 and Fcp1 contribute to maintain proper Rpb1-CTD-P levels, which have a role in transcription elongation and termination. Our findings show that Rpb4 phosphorylation plays an important function in RNAPII transcript biogenesis (mRNA and snoRNA), RNAPII stoichiometry, and thus likely in mRNA imprinting and buffering.

**Figure 1.**
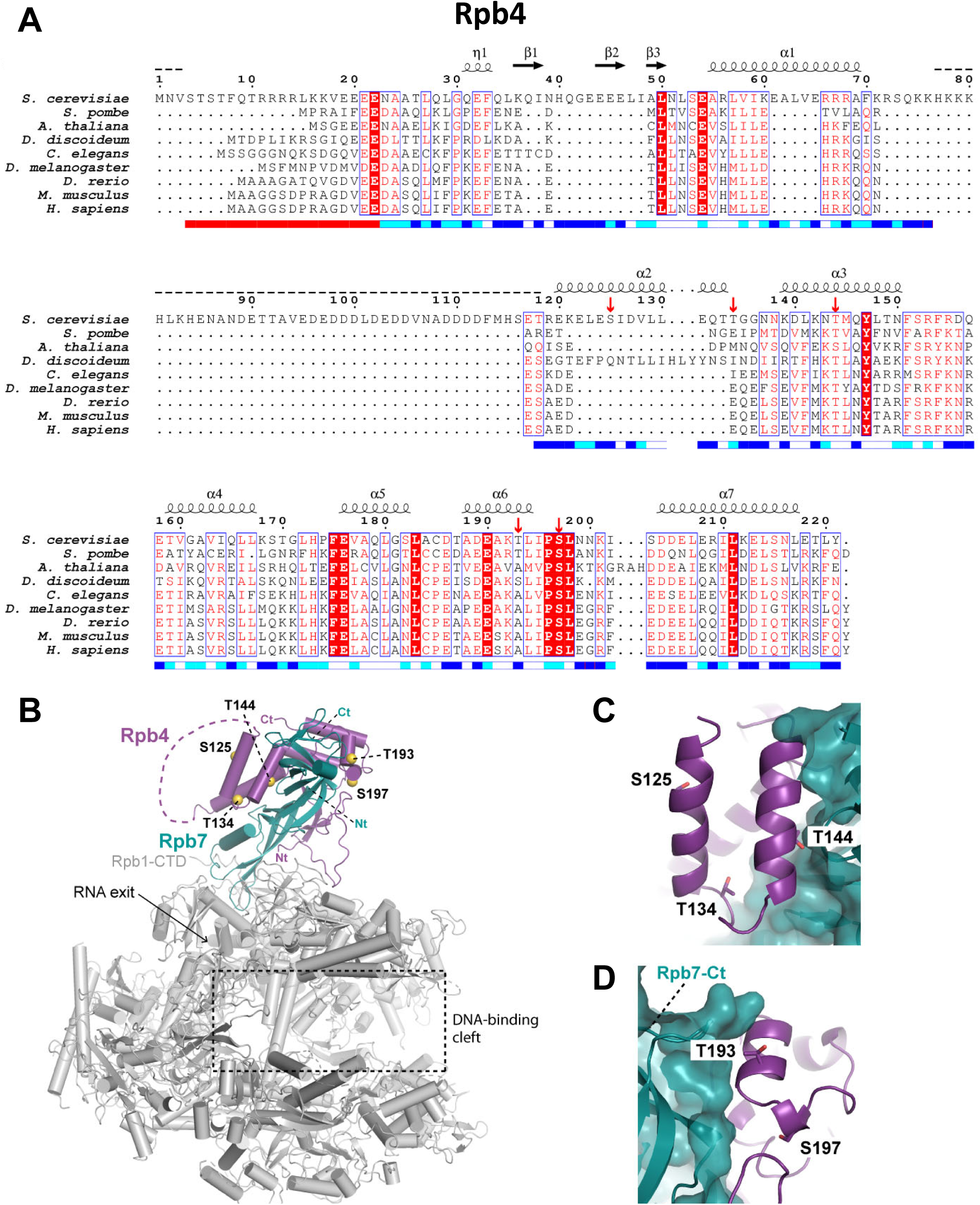
Structural location of potential Rpb4 phospho-sites. (A) Sequence alignment of Rpb4 from different species, performed with Clustal-Omega (Sievers et al. 2011) and represented with ESPript3 (Robert and Gouet 2014). Above the sequences, structural elements in yeast Rpb4 are shown as arrows for strands and springs for helices, while disordered regions are indicated with dashed lines. Below the sequences, solvent accessibility is shown as blue, cyan, and white boxes for accessible, partially accessible, and buried residues, respectively. Residues mutated in this study are marked with red arrow above the sequence. (B) The overall structure of yeast Pol II (PDB – 1WCM) (Armache et al. 2005); residues mutated in this study are shown as yellow spheres. Dashed lines correspond to disordered regions. (C) Close-up view of the region encompassing the residues S125, T134, and T144. (D) Close-up view of the region encompassing the residues T193 and S197.

## RESULTS

### Rpb4 is phosphorylated *in vivo*, and mutagenesis of Rpb4 phospho-sites causes growth and RNAPII integrity defects

Richard *et al*. (Richard et al. 2021) tandem affinity purified (TAP) Rpb4 from *S. cerevisiae* and analyzed its PTM by mass spectrometry (MS). They found that five residues are phosphorylated: two serines, S125 and S197, and three threonines, T134, T144, and T193. Sequence alignment of Rpb4 from several organisms from yeast to human (Fig. 1A) showed that S125 and T134 are exclusively present in *S. cerevisiae*, with S125 localized within a region, absent in other organisms, that is rich in charged amino acids. T193 is only present in *S. cerevisiae* and *Schizosaccharomyces pombe*, whereas at this position *Dictyostelium discoideum* presents a serine and other organisms harbor an alanine, a non-phosphorylatable residue. In contrast, T144 and S197 are highly conserved amino acids. S197 is present in all analyzed organisms within a highly conserved region (residues 189-198). T144 is only absent in *Arabidopsis thaliana*, which instead contains a serine that can also be phosphorylated. Fig. 1B shows the RNAPII structure from *S. cerevisiae* (Armache et al. 2005); identified phospho-sites in Rpb4 are highlighted. The structure shows that S125, T134, and T144 are located in a helix-loop-helix motif preceded by a disordered segment that is unique to *S. cerevisiae*, while T193 and S197 belong to a conserved helix-loop motif on the opposite side of Rpb7 (Fig. 1B-D). Notably, S125 and T193 are exposed to the solvent (Fig. 1A, C, and D), and thus readily accessible to modifying enzymes. Although T134 and S197 are partially accessible, they are localized at exposed loops that may undergo conformational changes (Fig. 1A, C, and D). In contrast, T144 is buried in the Rpb4/7 interface (Fig. 1A and C), suggesting that it may only be accessible upon Rpb7 detachment from Rpb4.

To study Rpb4 phosphorylation *in vivo*, we mutated all five of the aforementioned residues. We generated two mutants where the five phospho-sites were replaced either by alanine, a non-phosphorylatable residue, or aspartate, a negatively charged amino acid mimicking phosphorylation. The resulting mutants are termed *rpb4-S/T-A* (phospho-null) and *rpb4-S/T-D* (phospho-mimetic), respectively. We tagged Rpb4 at the C-terminal with the TAP epitope (Rpb4-TAP) in the two mutants and wild type (*wt*) cells for further complex purifications (Fig. S1A).

First, we analyzed whether these mutations cause defects in yeast cell growth. *RPB4* is a non-essential gene, but its deletion causes strong growth defects at 28° and 34°C, and thermo-sensitivity at 37°C (Fig. 2A) (Choder and Young 1993; Allepuz-Fuster et al. 2014). When all five phospho-residues were replaced with alanine (*rpb4-S/T-A*), cells grew slower than *wt* at all tested temperatures. Remarkably, when the five phospho-sites were replaced with aspartate (*rpb4-S/T-D*), the cells displayed strong growth defects and could barely grow at 37°C. The observation that mimicking a phosphorylated or unphosphorylated state of Rpb4 compromises cell growth suggests that both the phosphorylated and unphosphorylated states are important and that each of them functions transiently (see the Discussion).

**Figure 2.**
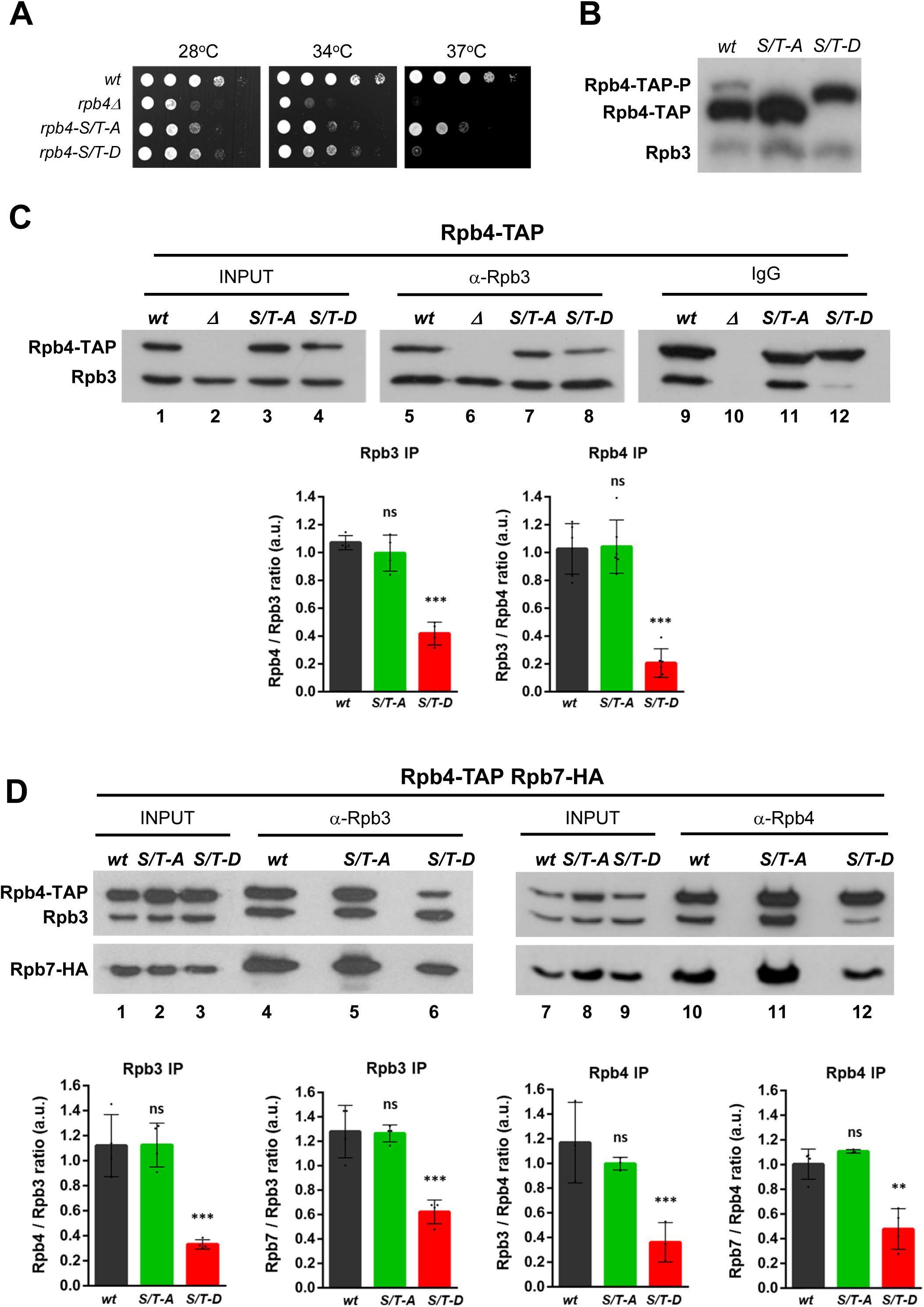
Analysis of the *rpb4* mutant phenotypes. (A) Cells were grown in rich media at the indicated temperatures for 2 days. (B) Analysis of Rpb4 phosphorylation in SDS-PAGE gels containing Phos-tag. Rpb3 was tested as a loading control. (C and D) Co-IP assays. WCE were used to immunoprecipitate RNAPII core with anti-Rpb3. Rpb4 was immunoprecipitated with either IgG Sepharose beads or anti-Rpb4, and Rpb7 with anti-HA. The levels of Rpb3, Rpb4 and Rpb7-HA were analyzed by western blot with the corresponding antibodies. The graphs in (C) show the Rpb4/Rpb3 and Rpb3/Rpb4 ratios, and the graphs in (D) show also the Rpb7/Rpb3 and Rpb7/Rpb4 ratios. All the ratios were calculated with values obtained by quantifying the corresponding band signals from at least three independent experiments.

We next analyzed whether Rpb4 is indeed phosphorylated *in vivo*. We prepared whole-cell extracts (WCE) from *wt*, *rpb4-S/T-A*, and *rpb4-S/T-D* cells; ran them in polyacrylamide SDS-PAGE gels containing Phos-tag (Nagy et al. 2018), and performed western blotting using an anti-Rpb4 antibody (Fig. 2B). The Phos-tag reagent can provoke a shift in the electrophoretic mobility of phosphorylated proteins and proteins containing negatively charged amino acids, such as aspartate. We detected two bands, of which the slow-migrating band corresponds to Rpb4-P. Importantly, Rpb4 phosphorylation was abolished in *rpb4-S/T-A*. The lack of the slowly migrating isoform in these cells suggests that the analyzed residues are likely accountable for most, if not all, of the Rpb4 phosphorylation events. It is also worth mentioning that only a minor portion of Rpb4 was phosphorylated in *wt* cells (Fig. 2B). This finding aligns with our suggestion that the phosphorylation is temporary; alternatively, only a small pool of *wt* Rpb4 undergoes constitutive phosphorylation. In *rpb4-S/T-D* cells, we only detected the slow-migrating band, mimicking hyperphosphorylation. Therefore, our results show that Rpb4 is a phosphoprotein and that the identified residues largely contribute to its phosphorylation *in vivo*.

We next examined RNAPII complex and Rpb4/7 heterodimer integrity in *wt*, *rpb4-S/T-A*, and *rpb4-S/T-D* cells by co-immunoprecipitation (Co-IP). We used anti-Rpb3 to immunoprecipitate RNAPII and anti-Rpb4 to immunoprecipitate the Rpb4/7 heterodimer (Fig. 2C). We included *rpb4Δ* cells as a mock control for Rpb4 IP. Rpb3 IP showed that in *rpb4-S/T-A* cells, where Rpb4 phosphorylation is abolished, the integrity of RNAPII remains unaffected because we detected similar levels of Rpb4-TAP compared with *wt* cells. This result suggests that phosphorylation of Rpb4 is not necessary for Rpb4/7 assembly into the RNAPII complex, but it might be required for subsequent steps during transcription. We confirmed this result after Rpb4-TAP purification and MS analysis of all RNAPII subunits (Fig. S1B). We next immunoprecipitated the core RNAPII using Rpb3 as the bait and observed that, despite the fact that comparable amounts of Rpb3 across all strains were immunoprecipitated (Fig. 2C, lanes 5-8), the levels of Rpb4-TAP were diminished in the *rpb4-S/T-D* cells compared with the *wt* and *rpb4-S/T-A* cells (Fig. 2C, see the difference between lanes 5 and 8, and see the Rpb4/Rpb3 ratio). On the other hand, when immunoprecipitating equivalent amounts of Rpb4 (Fig. 2C, lanes 9-12), the levels of Rpb3 remained unaltered in *rpb4-S/T-A*, but were significantly reduced in *rpb4-S/T-D* (Fig. 2C, compare lanes 9 and 12, and see the Rpb3/Rpb4 ratio in the graph).

We then C-terminally tagged Rpb7 with the 3xHA epitope in the three strains to analyze Rpb7 levels and to study Rpb4/7 heterodimer integrity. We noted similar immunoprecipitated amounts of Rpb3 in all strains (Fig. 2D, left panel, lanes 4-6), but Rpb4 and Rpb7-3xHA levels associated with the core RNAPII (Rpb3) were significantly reduced in *rpb4-S/T-D* compared with *wt* and *rpb4-S/T-A* cells (Fig. 2D, see the Rpb4/Rpb3 ratio in the graph). Consistent with this finding, when immunoprecipitating Rpb4 (Fig. 2D, lanes 10-12), levels of Rpb3 and Rpb7-3xHA were severely reduced in *rpb4-S/T-D*. Therefore, in this mutant, Rpb4/7 assembly to RNAPII as well as Rpb4/7 heterodimer formation are altered (Fig. 2D, see the Rpb7/3 and Rpb7/Rpb4 ratios in the graphs). The effects on RNAPII integrity appear stronger in cells expressing Rpb7-3xHA (compare the Rpb4/Rpb3 ratios in Fig. 2C and D), and, consistently, *rpb4-S/T-A* and *rpb4-S/T-D* cells expressing Rpb7-3xHA presented a stronger growth defect than untagged Rpb7 cells (Fig. S1C), specially *rpb4-S/T-D* mutant. The presence of 3xHA at the C-terminus of Rpb7 may weaken the Rpb4-Rpb7 association, as the Rpb7 C-terminus lies next to Rpb4, in the vicinity of phosphorylated residues (Fig. 1B and D).

The defect in RNAPII integrity could explain the growth defect of the *rpb4-S/T-D* cells, which, at least at 37 °C, closely resembles the phenotype of the *rpb4Δ* mutant. Indeed, *rpb4-S/T-D* cells hardly expressed Rpb4-TAP at 37°C (Fig. S1D). It is possible that the negative charge of aspartate residues impairs formation of a stable Rpb4/7 complex, in particular T144D, which is localized in a region that is important for the interaction between Rpb4 and Rpb7 (Fig. 1C and D). Hence, phosphorylation of Rpb4, and in particular residue T144, might impair heterodimer formation. It has been previously demonstrated that Rpb4 is important for the capacity of Rpb7 to bind RNAPII (Sheffer et al. 1999). Weakening the Rpb4-Rpb7 interaction, in combination with weak Rpb7 binding to RNAPII, can explain the reduced binding of the stalk to RNAPII in *rpb4-S/T-D* cells (Fig. 2C).

### Rpb4 phosphorylation plays a role in transcription regulation

To study the role of Rpb4-P in transcription, we used a simple method to analyze transcription elongation defects: the GLAM (<u>g</u>ene length-dependent <u>a</u>ccumulation of <u>m</u>RNA) assay that, when coupled to reverse transcription and quantitative real time PCR (RT-qPCR), is very useful to specifically analyze the transcription elongation efficiency depending on the length of the transcripts and independent of defects in initiation (Morillo-Huesca et al. 2006). As shown in Fig. S2A (left panel), the GLAM ratio (see the figure legend for details) was reduced in the two mutants, *rpb4-S/T-A* and *rpb4-S/T-D*, compared with *wt* cells, due to a reduction in the phosphatase activity expressed from the long transcripts, and as a consequence of defects in mRNA accumulation detected (Fig. S2A, right panel). We also tested the sensitivity of the cells to 6-azauracil (6AU), a drug often used to identify mutations affecting transcription elongation (Riles et al. 2004). Both *rpb4* phospho-mutants, *rpb4-S/T-A* and *rpb4-S/T-D* are resistant to 6AU at the permissive temperature of 28°C (Fig. S2B). The resistance of *wt* cells is due to the induction of the *IMD2* gene in conditions that lower GTP levels, as the treatment of 6AU does (Exinger and Lacroute 1992). However, *rpb4-S/T-A* cells are quite sensitive to 6AU at 34° and 37°C. Increasing the growth temperature and indeed the metabolism, the *de novo* synthesis of *IMD2* may be affected due to transcription elongation defects and therefore cause sensitivity to this drug.

We next studied the RNAPII association along the chromatin of several genes by chromatin immunoprecipitation (ChIP) followed by qPCR. We chromatin immunoprecipitated Rpb3 in *wt*, *rpb4-S/T-A*, and *rpb4-S/T-D* cells, and as a control we included *rpb4Δ* cells, which are known to have altered RNAPII chromatin association (Runner et al. 2008; Allepuz-Fuster et al. 2014). Thus, we analyzed Rpb3 association with the promoter (P), coding (CD) and 3′-end regions of four constitutively transcribed genes (*PMA1*, *YEF3*, *PGK1*, and *PYK1*; Fig. S3), which are highly expressed. In general, the effects on Rpb3 and Rpb4 occupancy were greater in *rpb4-S/T-D*, agreeing with the RNAPII assembly defects in that mutant. On average, the Rpb3 reduction was 20%-40% in *rpb4-S/T-A* cells and 20%-60% in *rpb4-S/T-D* cells. This result agrees with the elongation efficiency defects detected by the GLAM assay (Fig. S2A) and, in the particular case of the phospho-mimetic mutant, *rpb4-S/T-D*, with a defect in the association of Rpb4 with RNAPII. Moreover, these mutations also affected transcription initiation (see the “P” panels in Fig. S3).

Thereafter, we carried out genome-wide experiments in *wt* and *rpb4-S/T-A* cells. We did not include the phospho-mimetic mutant because it would be difficult to distinguish whether any detected defect in the additional steps during transcription is due to a defect in phosphorylation or altered RNAPII assembly. We first performed ChIP-seq to analyze Rpb3 and Rpb4 gene occupancy using chromatin from *wt* and *rpb4-S/T-A* cells. On average, Rpb3 occupancy at protein-coding genes was different in *rpb4-S/T-A* compared with *wt* cells (Fig. 3A). In *rpb4-S/T-A* cells, there was less Rpb3 associated at the transcription start site (TSS) and coding regions than in *wt* cells. However, this association pattern changed in the last third of the gene, where Rpb3 occupancy levels were similar to *wt* cells. Regarding Rpb4 occupancy, it was lower than in *wt* cells in the first half of the genes, but it clearly increased when reaching the transcription termination site (TES). In fact, Rpb4 association with chromatin was significantly higher in *rpb4-S/T-A* than in *wt* cells. Therefore, the ChIP-seq data suggest a defect in Rpb4 dissociation from RNAPII upon transcription termination. This is much clearer when we examine the Rpb4/Rpb3 ratio along the gene positions. As shown in Fig. 3B, in the second half of the genes, the Rpb4/Rpb3 occupancy ratio was higher in *rpb4-S/T-A* cells than in *wt* cells. When we represented the Rpb4/Rpb3 ratio in a heat map (Fig. 3C), where all *S. cerevisiae* genes are shown, we found that *wt* cells had genes with high ratios and genes with low ratios, but all genes in the phospho-mutant presented a similar Rpb4/Rpb3 association ratio. This result suggests that the lack of Rpb4 phosphorylation differentially affects Rpb4 association with RNAPII to transcribe different genes.

**Figure 3.**
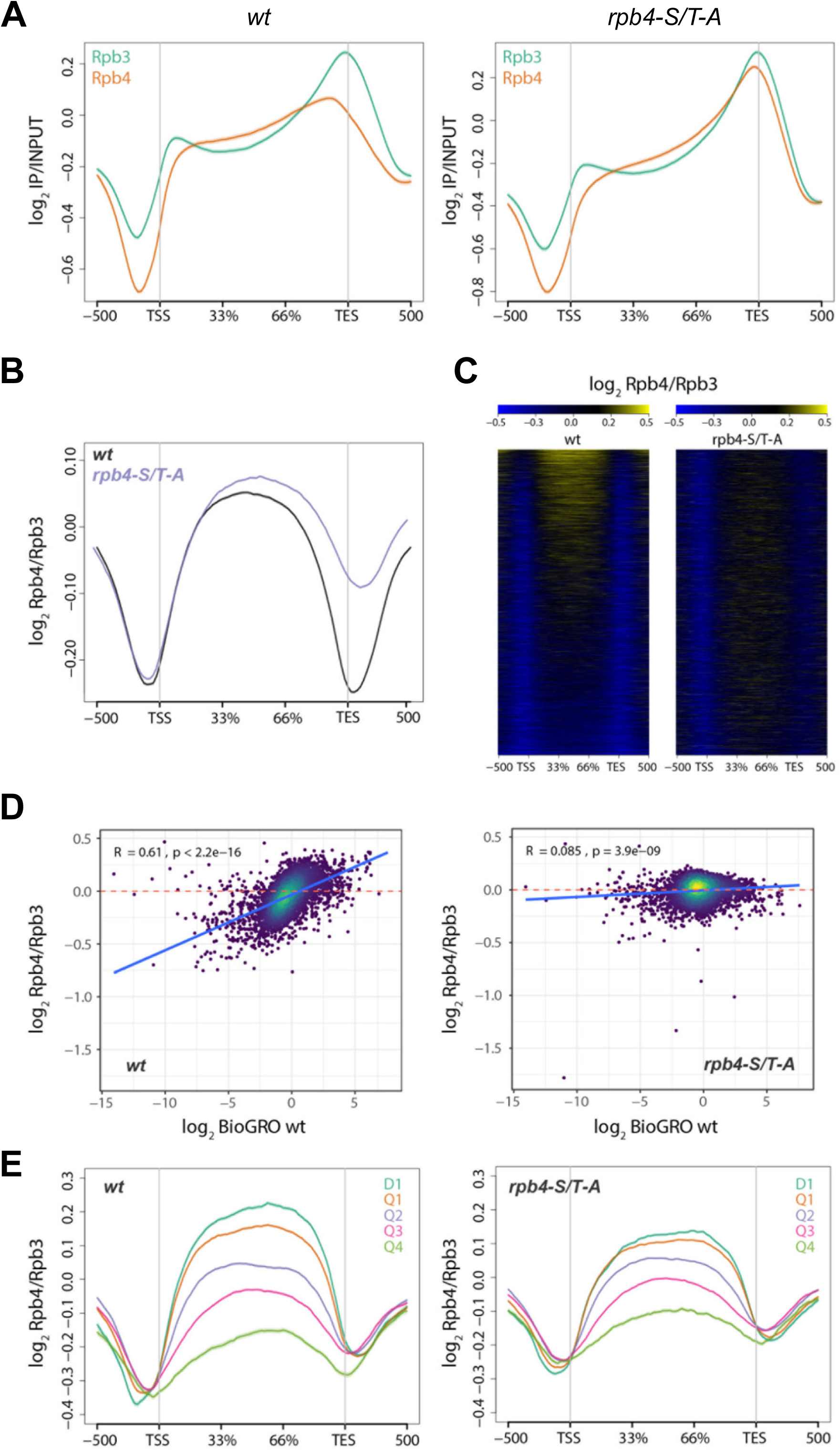
(A) Average metagene profile of input-normalized Rpb3 and Rpb4 ChIP-seq data over gene bodies and flanking regions of protein-coding genes (n = 5144) in *wt* (left) and *rpb4-S/T-A* (right) cells. (B) The Rpb4/Rpb3 ratio represented as the average metagene profile. (C) Associated heatmap representation of the Rpb4 signal relative to the Rpb3 signal over gene bodies and flanking regions of protein-coding genes (n = 5144). Rows on the heatmap represent individual genes. (D) Scatterplot showing genome-wide correlations between transcription rates, measured by BioGRO-seq (Begley et al. 2020), and Rpb4 enrichment, presented as the ratio of Rpb4/Rpb3 ChIP-seq signal, for all protein-coding genes. R values and their associated *p*-values correspond to Spearman correlations. Regions with a high density of overlapping genes are represented in yellowish tones in the color scale. (E) Average metagene plots of Rpb4/Rpb3 ChIP-seq signal separated according to five tiers of transcription strength. Q stands for quartile, and D for decile.

We also represented the genome-wide relationship between transcription rates, measured by BioGRO-seq (Begley et al. 2020), and Rpb4/Rpb3 ChIP-seq signal for all protein-coding genes. We demonstrated a direct correlation between the transcription level and Rpb4 enrichment in *wt* cells that is lost in the phospho-mutant, resulting in similar Rpb4 enrichment levels along the transcription rate range (Fig. 3D). To examine this effect in more detail, we performed metagene analysis of the Rpb4/Rpb3 signal in five subsets of genes according to the transcription rate (Fig. 3E). We observed that the positive correlation between Rpb4/Rpb3 and transcription rate in *wt* cells (Fig. 3D) was lost in the phospho-null mutant because the most expressed genes (D1 and Q1) lost the association with Rpb4, whereas the least expressed (Q2 and Q4) gained the association with Rpb4. We also noticed that the Rpb4/Rpb3 ratio was higher at the TSS, mostly for Q4 genes, although the general shape of the profiles did not change substantially. In summary, genes in the *rpb4-S/T-A* mutant lose Rpb4 proportionally to their level of transcription, so the most transcribed genes in *wt* cells are the ones that lose Rpb4 the most in the phospho-null mutant.

### Defects in Rpb4 phosphorylation change the levels of hundreds of mRNAs

Rpb4 is involved in mRNA decay (Lotan et al. 2005; Lotan et al. 2007; Goler-Baron et al. 2008), in addition to its function in transcription. Thus, Rpb4 is involved in determining the balance between mRNA synthesis and decay, a mechanism termed mRNA buffering, which is modulated by methylation of four Rpb4 residues: E19, E20, E21, and E22 (Richard et al. 2021). Here we examined whether the differences in chromatin association of Rpb4 with RNAPII also impacts mRNA abundance. To this end, we first analyzed by deep sequencing the transcriptomes of the *wt*, *rpb4Δ*, and *rpb4-S/T-A* strains. As a first approach to evaluate the similarity of the samples, we performed multidimensional scaling (MDS) analysis (Fig. S4A), which clearly separated the transcriptomes by strain, with *rpb4Δ* showing the most differences in the two first dimensions. Differential expression (DE) analysis revealed that a large fraction of the transcriptome was significantly altered in *rpb4Δ* cells compared with *wt* cells (Fig. 4A), consistent with a role for Rpb4 in mRNA buffering. Remarkably, *rpb4-S/T-A* cells also exhibited an altered transcriptome, although the changes were more modest. In line with the results of MDS analysis, *rpb4Δ* contained 2817 DE genes—1455 upregulated and 1362 downregulated—whereas *rpb4-S/T-A* cells had 1424 DE genes—711 upregulated and 713 downregulated. Gene Ontology (GO) enrichment analysis revealed that *rpb4Δ* and *rpb4-S/T-A* had reduced levels of mRNAs related to ribosomal biogenesis, translation, and RNA export and transport (Fig. 4B, right panel). However, both mutants presented different categories of upregulated genes involved in growth, energy production, and lipid metabolism (Fig. 4B, left panel).

**Figure 4.**
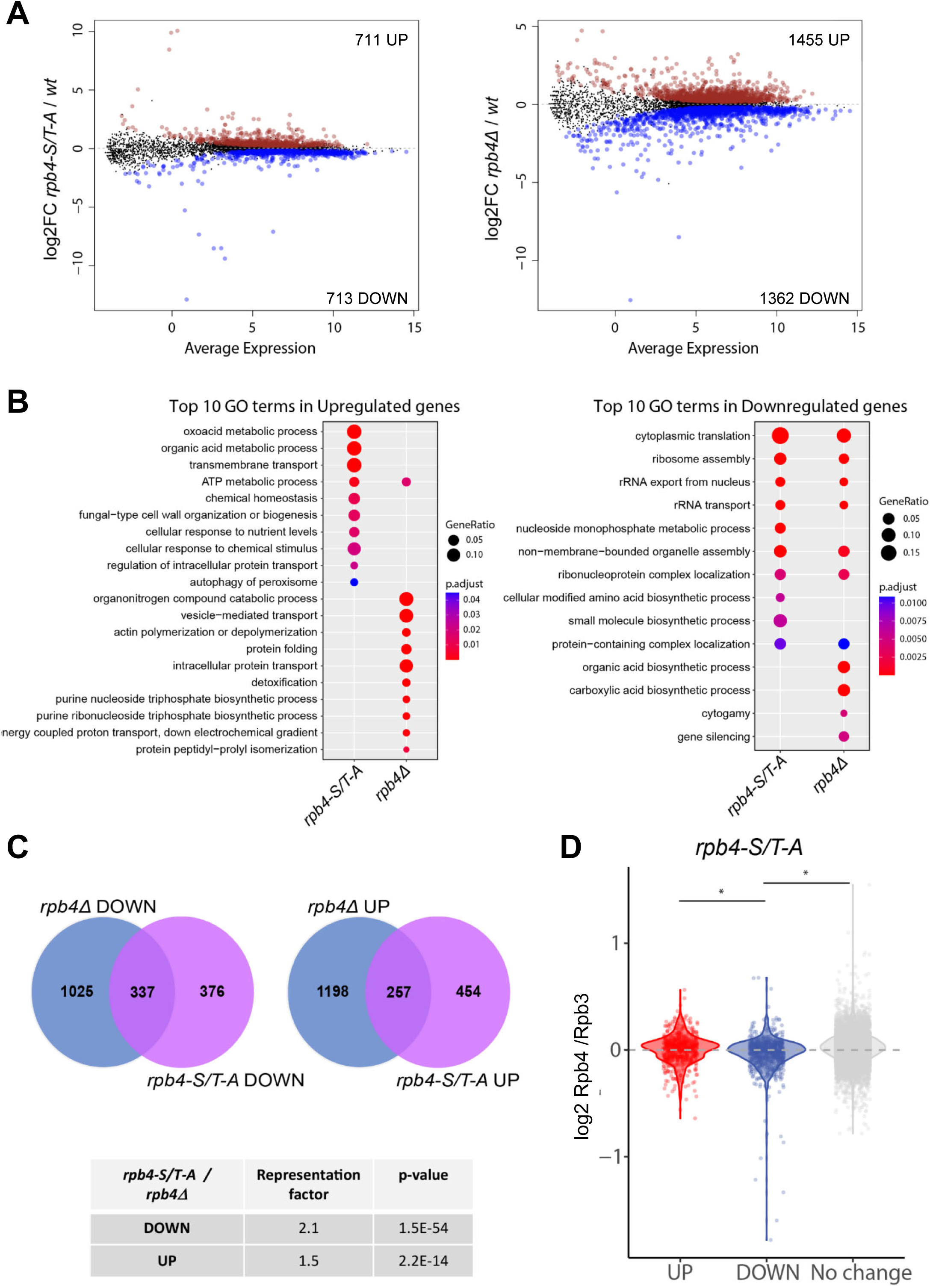
(A) MA plot of DE analysis results showing the genome-wide correlation between average mRNA levels and their changes in the two mutants compared with *wt* cells. Genes highlighted in red and blue are upregulated and downregulated, respectively (FDR < 0.05). (B) Comparative GO term enrichment in the lists of DE genes. The top 10 most significant GO terms enriched for up-and downregulated genes are shown. GeneRatio indicates the proportion of genes from the lists analyzed that have been mapped to specific GO terms. (C) Venn diagrams showing the overlap between DE gene sets from A. The *p*-values indicate the significance of the overlap, and representation factors > 1 indicate more overlap than expected by chance. (D) Boxplots with the Rpb4/Rpb3 ChIP-seq ratios in *rpb4-S/T-A* for the DE genes according to RNA-seq in *rpb4-S/T-A*/*wt*. Upregulated genes do not present higher relative Rpb4 occupancy. However, downregulated genes show a lower, but significant, relative occupancy of Rpb4 compared with upregulated and unchanged genes. \**p* ≤ 0.05.

We next sought to measure the similarities between the genes affected by each mutation. For this endeavor, we crossed the lists of DE genes (separating up-from downregulated genes) and found three distinct groups of genes in each list: (1) mRNAs that showed similar changes in the two mutants compared with *wt*, (2) mRNAs that changed only in the *rpb4Δ* mutant, and (3) mRNAs that changed exclusively in the *rpb4-S/T-A* mutant. The overrepresentation analysis results of mRNAs in group 1 was significant for up-and downregulated mRNAs, indicating that both mutations, *rpb4Δ* and *rpb4-S/T-A*, impact a roughly similar set of mRNAs (Fig. 4C).

To determine whether changes in mRNA levels are related to the association of Rpb4 with chromatin, we correlated the change in the Rpb4/Rpb3 ratio measured by ChIP-seq with the DE analysis results (Fig. 4D). In the vast majority of the genes, the Rpb4/Rpb3 ratio increased in the phospho-null mutant, but this outcome did not correlate with changes in mRNA levels, suggesting that mRNA buffering was not affected or that both transcription and mRNA decay were unaffected. Upregulated genes did not have a higher Rpb4/Rpb3 ratio than the genes whose expression did not change; therefore, changes in their expression cannot be explained only by an increase in Rpb4 association with them. Interestingly, we found that on average, downregulated genes had a lower Rpb4/Rpb3 ratio compared with non-DE genes in *rpb4-S/T-A* cells. This finding indicates that there is a correlation between low Rpb4 chromatin association and abnormally low mRNA levels.

In summary, transcriptome analysis indicates that Rpb4 phosphorylation is required to maintain proper expression of a significant fraction (∼30%) of yeast mRNAs. Moreover, downregulated genes, which account for 50% of total DE genes, have significantly less association with Rpb4. Indeed, these genes are also downregulated in the *rpb4Δ* mutant, supporting a direct role of Rpb4 and its phosphorylation in the proper expression of those genes.

### Fcp1 is required to maintain proper Rpb4-P levels

The function of Rpb4 is related to several CTD phosphatases (Allepuz-Fuster et al. 2014). Therefore, we tested whether Rpb4-P levels depend on the phosphatases that differentially dephosphorylate the Rpb1-CTD during the transcription cycle: Rtr1, Ssu72, Fcp1, and Glc7 (Kobor et al. 1999; Krishnamurthy et al. 2004; Mosley et al. 2009; Schreieck et al. 2014). We analyzed Rpb4-P levels using WCE from the *rtr1Δ*, *ssu72-2*, *fcp1-5001*, and *glc7-12* mutant strains, and *wt* cells (Fig. 5A). Interestingly, only the *fcp1-5001* mutant presented an increase in the intensity of the slower migrating band as a consequence of Rpb4 hyperphosphorylation, indicating that the activity of Fcp1 is required to maintain proper Rpb4-P levels. We calculated the Rpb4-P/Rpb4 ratio in *wt* cells and each of the phosphatase mutants. As shown in Fig. 5A (right panel), the *fcp1-5001* mutant had Rpb4-P levels that were at least three times higher than in *wt* cells, whereas the rest of the phosphatase mutants had levels similar to that of *wt* cells. Fcp1 is a Ser2P phosphatase whose function is important upon transcription termination to recycle RNAPII and to reinitiate a new round of transcription (Kobor et al. 1999; Cho et al. 2001). Fcp1 also dephosphorylates Thr4P in yeast (Allepuz-Fuster et al. 2014) and vertebrates (Hsin et al. 2014). A physical interaction between Rpb4 and Fcp1 has been reported in fission yeast and flies (Kimura et al. 2002; Kamenski et al. 2004; Tombacz et al. 2009). Moreover, we have previously demonstrated that Rpb4 associates with Fcp1 and Sub1 in the proper context of RNAPII to modulate Rpb1-CTD dephosphorylation (Allepuz-Fuster et al. 2014; Garavis et al. 2017).

**Figure 5.**
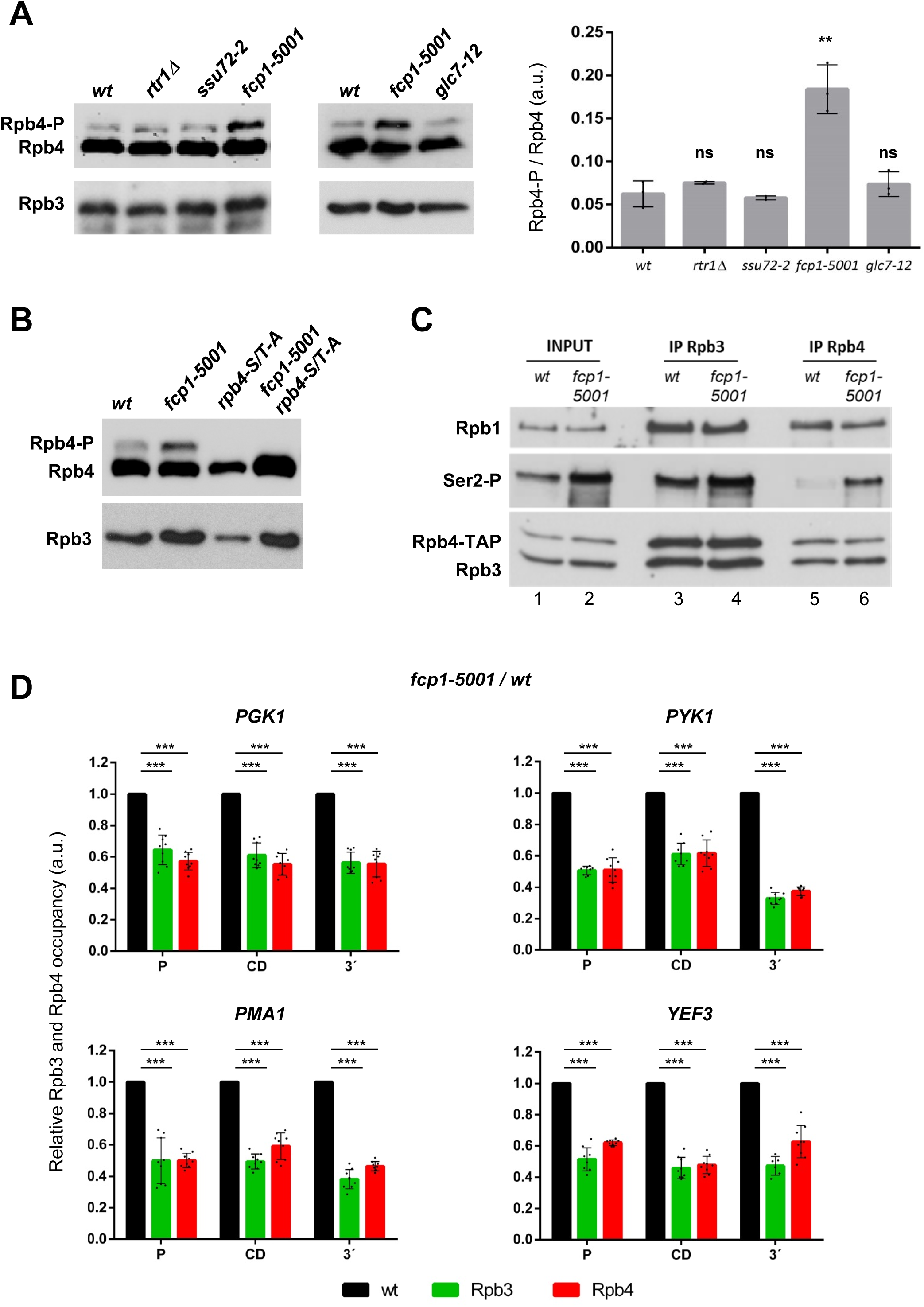
(A) The Rpb4-P level was specifically increased in the phosphatase mutant, *fcp1-5001*. Left, WCE from *wt* and the CTD phosphatase mutants *rtr1Δ*, *ssu72-2*, and *fcp1-5001* were run in SDS-PAGE gels containing Phos-tag and submitted to western blotting with anti-Rpb4 to examine levels of Rpb4-P and non-phosphorylated Rpb4. Right, plots of the Rpb4-P/Rpb4 ratio determined by quantitation (by densitometry) of immunoreactive bands from the western blots. The mean values and standard errors are from three independent experiments. ns, not significant; \*\**p* ≤ 0.01. (B) The Rpb4-P level is due to the phosphorylation of all, some, or any of the five phospho-residues mutated in *rpb4-S/T-A*. Analysis of the Rpb4-P level in *wt*, *fcp1-5001*, *rpb4-S/T-A*, and *fcp1-5001 rpb4-S/T-A* double mutant cells as in (A). (C) Co-IP assays to analyze RNAPII integrity in *wt* and *fcp1-5001* cells. WCE from *wt* and *fcp1-5001* were used to immunoprecipitate either Rpb3 or Rpb4 using the corresponding antibodies. Immunoprecipitated samples were tested by western blot to detect Rpb1, Rpb1-Ser2P, Rpb4-TAP, and Rpb3. (D) ChIP-qPCR assays to evaluate Rpb3 and Rpb4 association with the promoter (P), coding (CD), and 3′-end (3′) regions of several constitutively transcribed genes. The black bars represent *wt* levels set up as 1 for both Rpb3 and Rpb4 association. Anti-Rpb3 or anti-Rpb4 were used to immunoprecipitate both RNAPII subunits and immunoprecipitated DNA was tested by qPCR. \*\*\**p* ≤ 0.001.

We next aimed to determine whether residues other than the five identified phospho-residues could be the target of Fcp1 phosphatase activity, because three more Rpb4 phospho-sites were described in a phospho-proteomic study: S53, S203, and S215 (Lanz et al. 2021). While the S53 sidechain is buried, those of S203 and S215 are exposed on the surface of Rpb4/7 heterodimer and, therefore, could be a target of PTM (Gonzalez-Jimenez et al. 2021). We constructed the *rpb4-S/T-A fcp1-5001* double mutant. We reasoned that if some other Rpb4 residues are targeted by Fcp1, we should still expect to detect Rpb4-P in Phos-tag gels. However, this was not the case, and we did not detect any Rpb4-P isoforms in *rpb4-S/T-A fcp1-5001* cells (Fig. 5B), indicating that all or some of the five identified residues are indeed phosphorylated *in vivo*, and their dephosphorylation depends on functional Fcp1. If there are additional phospho-residues, they do not seem to be a target of Fcp1, and they might represent a minor fraction with respect to the total Rpb4-P and are undetectable in our assays (see *rpb4-S/T-A* lanes). We next performed Co-IP assays to evaluate whether RNAPII assembly is affected in the *fcp1-5001* mutant as a consequence of altered Rpb4-P, as we observed in the *rpb4-S/T-D* phospho-mimetic mutant. We immunoprecipitated Rpb3, and analyzed Rpb4, Rpb1 and Rpb1-CTDSer2P levels. In agreement with Fcp1 phosphatase specificity (Kobor et al. 1999; Mandal et al. 2002), we detected Ser2P hyperphosphorylation in *fcp1-5001* cells (Fig. 5C), although similar levels of Rpb4 were assembled into RNAPII. This was quite significant in the case of Rpb4 IP (compare lanes 5 and 6). Curiously, we did not detect any assembly defect, suggesting that likely not all five phospho-sites are targets of Fcp1, as their phosphorylation would contribute to stalk dissociation.

We next studied Rpb3 and Rpb4 occupancy levels in *wt* and *fcp1-5001* cells in several constitutively transcribed genes and observed a significant reduction of RNAPII association with genes from the promoter region to the 3′-end (Fig. 5D). Fcp1 is involved in the elongation step (Mandal et al. 2002; Kong et al. 2005) and, therefore, the ChIP data do not exclude that reduced Rpb3 levels could provoke defects in elongation that are not only due to increased Ser2P and Rpb4-P levels. In the *fcp1-5001* mutant, there was reduced Rpb4 association with chromatin. Because Fcp1 does not affect the Rpb4-RNAPII interaction, this finding suggests that low association of Rpb4 with chromatin is due to a direct effect of *fcp1-5001* on chromatin binding of RNAPII. In summary, our data show that some or all of the identified phospho-residues contribute to Rpb4 phosphorylation and that a functional Fcp1 is needed to maintain proper Rpb4-P levels. The results again suggest a role for Rpb4-P levels in transcription termination/RNAPII recycling.

### Hrr25 specifically phosphorylates Rpb4

To identify the kinase(s) targeting Rpb4, we expressed a recombinant heterodimer Rpb4/Rpb7-6xHIS (Garavis et al. 2017) and incubated it with WCE from *rpb4Δ* cells in a pull-down assay. We used WCE from *rpb4Δ* cells because, in the absence of *RPB4*, Rpb7 levels are quite reduced and can hardly be detected (Edwards et al. 1991; Sheffer et al. 1999). In this manner, we forced the interactions with rRpb4/Rpb7. Thereafter we performed MS analysis to identify Rpb4/7-interacting peptides. We focused on nuclear kinases and found that Kin28 and Bur1, two canonical Rpb1-CTD kinases (Qiu et al. 2009), and Hrr25, a non-canonical CTD kinase (Nemec et al. 2019), interact with Rpb4/7 (Fig. S5A). Kin28 is known to phosphorylate Ser5 and Ser7 within the CTD repetitions during transcription initiation (Komarnitsky et al. 2000; Buratowski 2003; Glover-Cutter et al. 2009). Bur1 phosphorylates Ser2 near the promoter regions, during early elongation (Qiu et al. 2009), Ser7 to promote elongation and to suppress cryptic transcription (Tietjen et al. 2010), and more recently it has been shown to phosphorylate Thr4 (Nemec et al. 2019). Moreover, it can also phosphorylate Ser5P *in vitro* (Garcia et al. 2010). Hrr25 phosphorylates Thr4 in a gene-class-specific manner and is involved in transcription termination (Nemec et al. 2019). We corroborated by CoIP the interaction between Rpb4 and Hrr25 (Fig. 6A) as well as Kin28 and Bur1 (Fig. S5B). The interaction with the last two kinases is not surprising because it is well known that they both target Rpb1-CTD (Rodriguez et al. 2000; Qiu et al. 2009). Therefore, all of them have been shown to interact with the transcriptional machinery. However, we did not find any other CTD kinase, such as Srb10 or Ctk1, also known to phosphorylate Rpb1-CTD.

**Figure 6.**
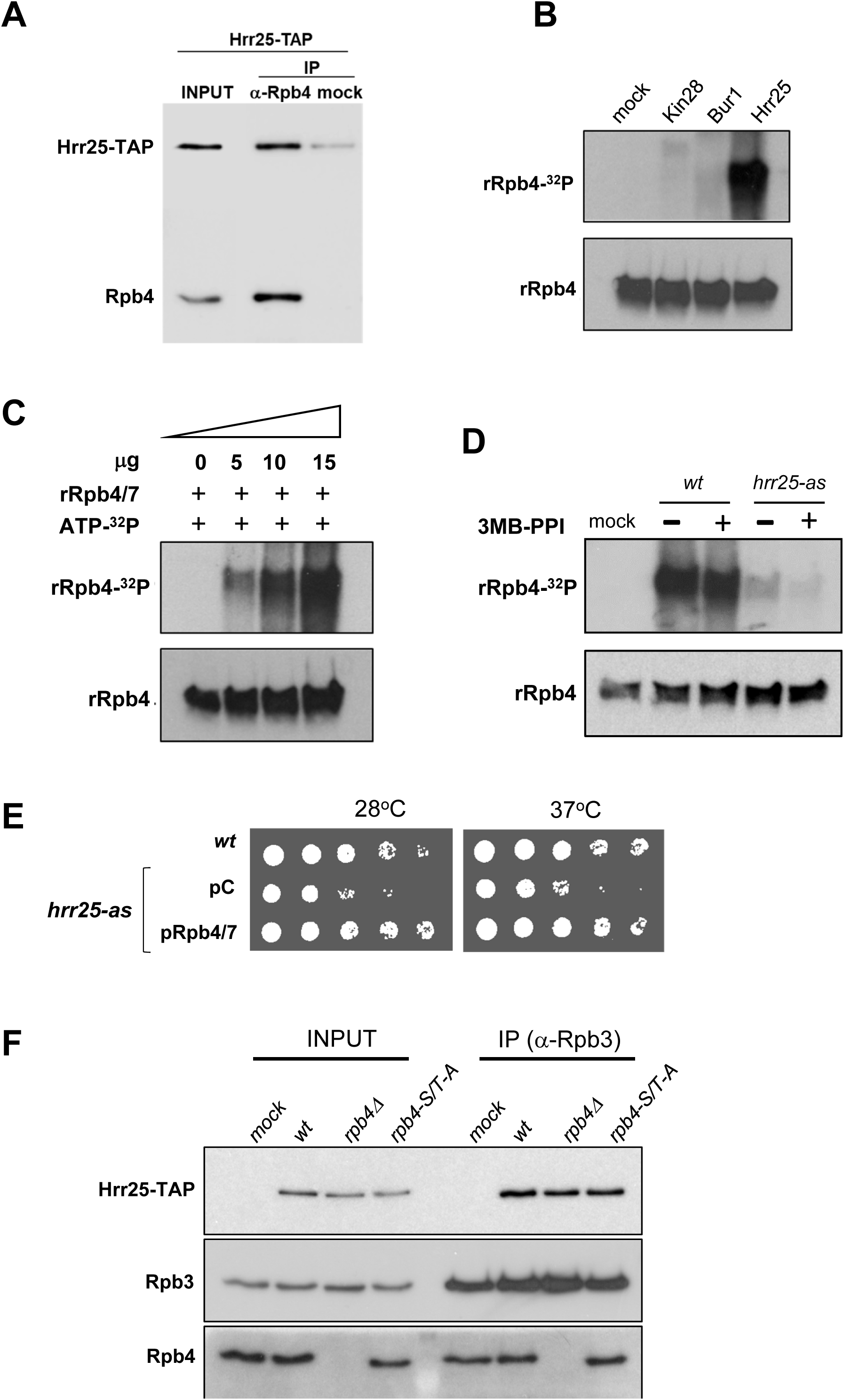
Hrr25-TAP interacts with Rpb4. (A) WCE from the Hrr25-TAP strain was used to immunoprecipitate Rpb4 with the anti-Rpb4 antibody coupled to mouse Dynabeads. To detect nonspecific binding, WCE was incubated with Dynabeads but without antibody (mock). The input and immunoprecipitated samples were submitted to western blotting to analyze the level of Hrr25-TAP associated with Rpb4. (B) *In vitro* kinase assays were performed to phosphorylate Rpb4 using the indicated purified kinases, [γ-^32^P]-ATP, and rRpb4/7-6XHis as a substrate. The reactions were loaded in polyacrylamide gels and transferred to a nylon membrane that was first exposed to detect radioactive rRpb4-^32^P (Rpb4-P, upper panel), and then submitted to western blotting with an anti-Rpb4 antibody (rRpb4, bottom panel) as a control to ensure that similar amounts of Rpb4 protein were used in each reaction. (C) *In vitro* kinase assays as in (B), but using increasing amounts of purified Hrr25. (D). *In vitro* kinase with purified Hrr25 from *wt* and the *hrr25-as* cells expressing Hrr25-TAP grown in the presence of the kinase inhibitor 3MB-PPI (+) or DMSO (-). (E) Overexpression of *RPB4/7* partially rescues *hrr25-as* growth phenotypes. Strains with the indicated genotypes and transformed with an empty plasmid (pC) or co-expressing *RPB4/7* were grown in selective minimal medium at the indicated temperatures for 2 days. (F) Co-IP and western blotting to analyze the interaction between Hrr25-TAP and RNAPII, dependent on Rpb4, in *wt*, *rpb4Δ*, and *rpb4-S/T-A* cells. RNAPII was immunoprecipitated through Rpb3, using an anti-Rpb3 monoclonal antibody. The Rpb3, Rpb4, and Hrr25-TAP levels were analyzed using the corresponding antibodies. WCE from a non-TAP tagged strain was used as a negative control (mock).

We next evaluated whether any of the three kinases could phosphorylate Rpb4 in an *in vitro* kinase assay performed with affinity-purified Kin28, Bur1, and Hrr25 (see Materials and methods) and radiolabeled ^32^P-ATP. First, we verified that the purified kinases were active using recombinant GST-CTD as substrate and testing for CTD-Ser5P in the case of Kin28 and Bur1, and CTD-Thr4P in the case of Hrr25. Interestingly, although the three kinases phosphorylated the CTD (Fig. S5C), only Hrr25 was able to phosphorylate Rpb4 (Fig. 6B). Increasing the amount of Hrr25 correlated with increased Rpb4 phosphorylation (Fig. 6C).

We took advantage of an analog-sensitive kinase mutant, *hrr25-as* (Nemec et al. 2019), in which a conserved residue in the ATP binding pocket has been substituted by glycine (I82G), allowing ATP analogs, such as 3MB-PPI to selectively dock to and inhibit the kinase activity non-covalently. We then constructed the *hrr25-as-TAP* strain to study the specific phosphorylation of Rpb4 by Hrr25. We purified *wt* and *hrr25-as* strains grown with or without 3MB-PPI as in a previous study (Nemec et al. 2019). We verified that *wt* and *hrr25*-as strains expressed a similar amount of Hrr25-TAP by analyzing WCE, and that comparable amounts of Hrr25-TAP were IgG precipitated (Fig. S6A, left panel) and purified (Fig. S6A, right panel). As shown in Fig. 6D, Hrr25 from *wt* strain phosphorylated Rpb4 in both conditions, whereas the *hrr25-as* mutant hardly phosphorylated it even in the absence of the drug, when the kinase activity should be similar to that of *wt* cells (Nemec et al. 2019). We observed a similar result in the case of CTD phosphorylation (Fig. S6B and C). Then, we tested for growth, and we observed that in the absence of the kinase inhibitor (DMSO, the vehicle of 3MB-PPI, Fig. S6D), the *hrr25-as-TAP* strain grew worse than the isogenic *hrr25-as* strain, which grew similar to *wt* cells. *hrr25-as-TAP* is probably not fully active when expressed with a TAP epitope, affecting its functionality even in the absence of the drug. Nonetheless, as expected both *hrr25-as* strains showed strong sensitivity to the kinase inhibitor as expected, consistent with the fact that *HRR25* is an essential gene (3MB-PPI, Fig. S6D and (Nemec et al. 2019). Our data show that Hrr25 phosphorylates Rpb4 and that a nonfunctional Hrr25 kinase can hardly phosphorylate it. Furthermore, and consistent with this result, overexpression of Rpb4/7 suppresses *hrr25-as* growth phenotypes (Fig. 6E), indicating that Rpb4 is a main target of Hrr25 activity. We performed Co-IP assays that showed Hrr25 association with RNAPII was similar in *wt*, *rpb4Δ*, and *rpb4-S/T-A* cells and, therefore, it does not depend on Rpb4 (Fig. 6E).

We also tested whether other known CTD kinases, such as Srb10 and Ctk1, phosphorylate Rpb4, and analyzed WCE from the *srb10Δ* and *ctk1Δ* mutants in Phos-tag gels. The *srb10Δ* and *ctk1Δ* mutants had a similar Rpb4 phosphorylation pattern to that of *wt* cells (Fig. S6E). This finding is consistent with the lack of Ctk1 and Srb10 among the rRpb4/7-interacting proteins in our pull-down assay coupled with MS. Therefore, we conclude that Hrr25 specifically phosphorylates Rpb4, as one of its main targets, but not any of the canonical Rpb1-CTD kinases that act during the transcription cycle.

### Rpb4 phosphorylation is required for snoRNA transcription termination and poly(A) site selection

We next investigated whether, as in the case of *rpb4-S/T-A* cells, the Rpb4/Rpb3 chromatin association ratio is altered by using the *hrr25-is* strain, which expresses an “irreversibly sensitized” (denoted by “-is”) kinase. In this mutant, two positions in the ATP pocket are altered, I82G and I23C; these mutations permit an irreversible and covalent inhibition of the Hrr25-is kinase by specific drugs (Nemec et al. 2019). We first tested the *wt* and *hrr25-is* growth phenotype in the presence of CMK that specifically inhibits the Hrr25 kinase activity of this mutant (Nemec et al. 2019). *hrr25-is* growth was similar to that of *wt* cells in DMSO (CMK vehicle)-containing rich media; however, its growth was significantly reduced when treated with CMK (Fig. S6F). We then performed ChIP-qPCR to analyze Rpb3 and Rpb4 gene occupancy in *wt* and *hrr25-is* cells treated with DMSO or CMK, and we calculated Rpb4/Rpb3 ratio (Fig. 7A, and see Materials and methods for details). Interestingly, we observed that the Rpb4/Rpb3 association ratio was significantly increased in the absence of a functional Hrr25 kinase, which agrees with our findings that Hrr25 is an Rpb4 kinase and that phosphorylation of Rpb4 is required for proper chromatin occupancy as we observed in the *rpb4-S/T-A* phospho-null mutant (Fig. 3B). Because transcription of small non-coding genes has been reported to be modulated by Hrr25 (Nemec et al. 2019), we also analyzed the Rpb4/Rpb3 ratio association with the *SNR13* and *SNR33* genes. We found that mutating *HRR25* compromises Rpb4 release from RNAPII engaged in snoRNA transcription (Fig. 7B). Moreover, our RNA-seq data unveiled that the genome-wide abundance of snRNAs was decreased in *rpb4-S/T-A* cells compared with *wt* cells (Fig. 7C).

**Figure 7.**
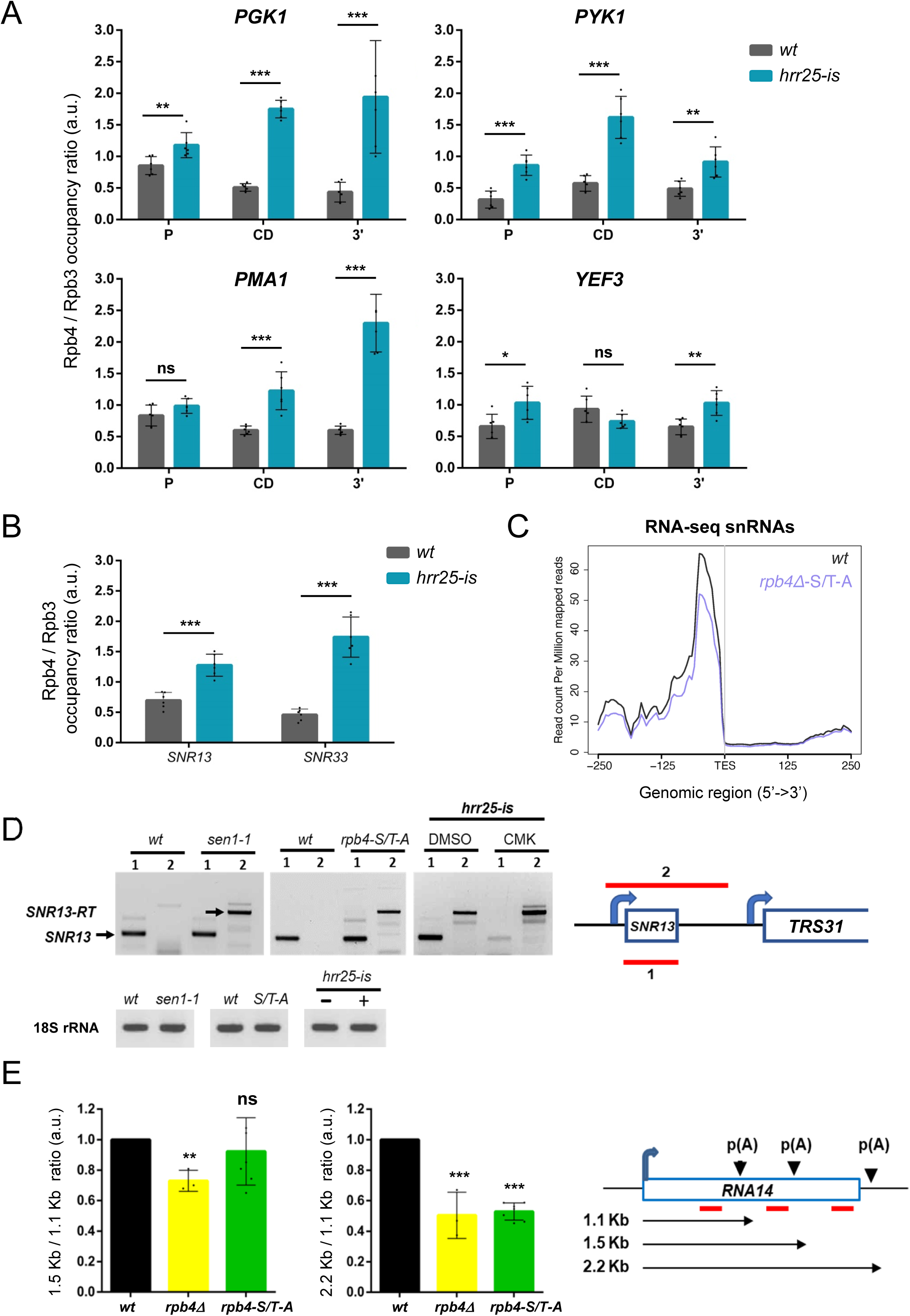
(A and B) The Rpb4/Rpb3 occupancy ratio. Rpb4 and Rpb3 ChIP was performed with cells grown in the presence of CMK or DMSO. Thereafter, Rpb4 and Rpb3 occupancy was examined by qPCR. The values obtained were processed as follow: First, the CMK/DMSO ratio was calculated independently for Rpb4 and Rpb3, and then the Rpb4/Rpb3 ratio was estimated and graphed. (A) shows the results for the protein-coding genes and (B) shows the results for the *SNR13* and *SNR33* genes, which encode two snoRNAs. (C) Polyadenylation-signal-aligned average metagene plot showing RNA-seq reads originating from snRNA genes in *wt* and mutant cells. (D) RT-PCR to analyze transcription termination defects of *snR13*. Total RNA was purified from the following strains: *wt* and *sen1-1* grown at 37°C; and *wt*, *rpb4-S/T-A*, and *hrr25-is* grown at 28°C. *hrr25-is* was grown in the presence of either DMSO or CMK. Full-length snR13 and readthrough (*snR13*-RT) transcripts are indicated by arrows. The level of 18S rRNA was used as a control. A schematic representation of the *SNR13* gene and 3′-end flanking region is shown, where the PCR products corresponding to mature snR13 (123 bp) or snR13-RT (419 bp) are represented by red bars. (E) RT-qPCR to analyze 3′-end processing in *wt*, *rpb4Δ*, and *rpb4-S/T-A* cells. Long and short *RNA14* transcripts were analyzed by qPCR, and the ratio of the obtained values (2.2 kb/1.1 kb and 1.5 kb/1.1kb) from at least three independent experiments were calculated. Then, the resulting data from the *rpb4Δ* and *rpb4-S/T-A* mutants relative to *wt* cells, whose values were set up to 1, were graphed. \**p* ≤ 0.05; \*\**p* ≤ 0.01; \*\*\**p* ≤ 0.001; ns, not significant. A schematic representation of the *RNA14* gene, poly(A) sites, and transcripts resulting from 3′-end processing is shown.

It has been reported a significant enrichment of Hrr25 at the 3ʹ-ends of snoRNAs and also throughout the length of protein-coding genes. Indeed, inhibition of Hrr25 kinase activity provokes termination defects of snoRNAs and elongation defects of protein-coding genes (Nemec et al. 2019). These authors proposed that Hrr25 phosphorylates the CTD in a gene-specific manner. Thus, we tested whether Rpb4 phosphorylation deficiency might affect transcription termination of snoRNAs. Specifically, we examined the well-known *SNR13* transcript (Kim et al. 2006; Franco-Echevarria et al. 2017) by using RT-PCR (Fig. 7D). As a positive control of snoRNA transcription termination defects, we used the *sen1–1* mutant grown at 37°C (Kim et al. 2006; Franco-Echevarria et al. 2017). We detected a strong termination defect of *SNR13* in *rpb4-S/T-A* and *hrr25-is* strains, similar to that of the *sen1-1* mutant (Fig. 7D). These results indicate that Rpb4 phosphorylation by Hrr25 is required for snoRNA transcription termination and are consistent with the reduction in snoRNAs levels detected by RNA-seq (Fig. 7C).

A previous study showed that the lack of Rpb4 alters polyadenylation site usage at the *RNA14* gene; the authors proposed a role for Rpb4 in coupling transcription and 3′-end processing of pre-mRNAs (Runner et al. 2008). Transcription of the *RNA14* gene generates three transcripts of 1.1, 1.5, and 2.2 kilobase pairs (kb) (Fig. 7E), as a result of the selection of three different poly(A) sites during 3′-end processing (Sparks and Dieckmann 1998). We tested whether Rpb4 phosphorylation deficiency could also affect polyadenylation site usage. For that purpose, we measured levels of *RNA14* long (2.2 and 1.5 kb) and short (1.1 kb) transcripts in *wt* and *rpb4-S/T-A* cells by RT-qPCR, and calculated the ratio of long versus short transcripts (Fig. 7E). We included in our assays *rpb4Δ* cells as a positive control for defects in co-transcriptional polyadenylation. In these cells, we detected an increase in the shortest transcript and a decrease in the longest ones (1.5 and 2.2 kb), as a consequence of the preference in the selection of the first poly(A) site, as described previously (Runner et al. 2008). In the *rpb4-S/T-A* phospho-null mutant, there was a significant reduction in the selection of the most distant and major poly(A) site, as in *rpb4Δ* cells (Fig. 7E, see the 2.2 kb/1.1 kb ratio graph), indicating that proper phosphorylation of Rpb4 is relevant for polyadenylation site usage (see Discussion).

We next evaluated whether Rpb4 phosphorylation deficiency causes similar effects as the absence of Hrr25 kinase activity, and explored whether *rpb4Δ* and *rpb4-S/T-A* mutants share transcriptomic defects with the catalytic *hrr25-is* mutant. For that purpose, we extracted RNA-seq data from (Nemec et al. 2019) and compared it with our RNA-seq data (Fig. 8; see Materials and methods for details). First, we evaluated the similarity of *wt* (control) and *hrr25-is* RNA samples by performing principal component analysis (PCA, Fig. 8A, left panel), which separated the transcriptomes by strain. DE analysis revealed that the *hrr25-is* mutant transcriptome was significantly altered compared with the *wt* transcriptome (Fig. 8A, right panel). *hrr25-is* contained 763 DE genes, 330 upregulated and 433 downregulated. We measured the similarities among the genes affected by the *rpb4Δ*, *rpb4-S/T-A*, and *hrr25-is* mutations by crossing the lists of DE genes (Fig. 8B). We found that: (1) 252 genes changed simultaneously in the three mutants compared with *wt*, which represent 33% of the total genes altered in the *hrr25-is* mutant; (2) the *rpb4-S/T-A* and *hrr25-is* mutants shared 318 DE genes (43%, p = 1.3e-32); (3) the *rpb4Δ* and *hrr25-is* mutants shared 236 DE genes (57%, p = 3.2e-10); (4) there were 1726 DE genes only in the *rpb4Δ* mutant; (5) there were 455 DE genes only in the *rpb4-S/T-A* mutant; and (6) there were 259 DE genes only in the *hrr25-is* mutant. Analysis of GO term enrichment for groups 1 (252 genes) and 2 (318 genes) showed that among the top 10 categories are translation, ribosomal assembly, ribosomal RNA (rRNA) export and transport (Fig. 8C). Consistently, the absence of Rpb4 phosphorylation provoked sensitivity to neomycin, a drug used to detect translation defects (Arenz and Wilson 2016), as it does the deletion of *RPB4, hrr25-is*, and *hrr25-as* mutations (Fig. 8D). Moreover, *RPB4/7* overexpression partially restored *hrr25-as* growth in the presence of neomycin (Fig. 8D), which again indicates that the *hrr25* mutant phenotypes are mostly due to the inefficient phosphorylation of Rpb4.

**Figure 8.**
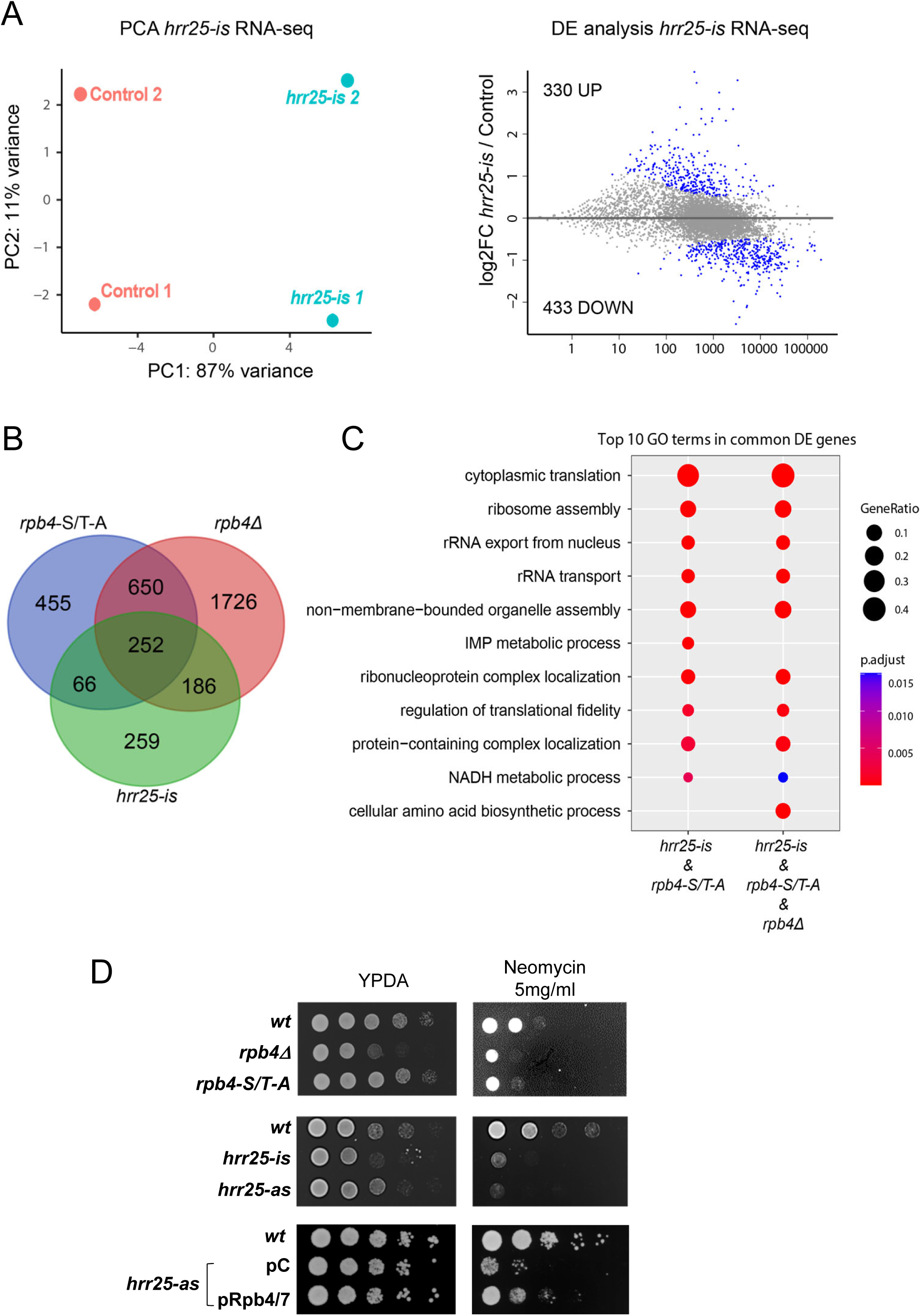
(A) PCA plot (left) and MA plot (right) from *wt* and *hrr25-is* RNA-seq samples (original datasets obtained previously (Nemec et al. 2019) and re-analyzed for this study). DE genes according to an FDR < 0.05. (B) Overlap between DE genes from *hrr15-is*, and from *rpb4Δ* and *rpb4-S/T-A* mutants (data from our RNA-seq experiments). Thirty-three percent of the genes altered in the *hrr25-is* mutant were also altered in the *rpb4Δ* and *rpb4-S/T-A* mutants (252 genes); 57% of them were shared with the *rpb4Δ* mutant (438 genes) and 43% were shared with the *rpb4-S/T-A* mutant (318 genes). (C) Comparative GO enrichment analysis dot plot showing the top 10 most significantly enriched GO terms associated with DE genes overlapping in the *hrr25-is* and *rpb4-S/T-A* mutants (318 genes total; representation factor 1.8 and *p* < 1.306e-32) and in the *hrr25-is* and *rpb4Δ* mutants (438 genes total; representation factor 1.22 and *p* < 3.16e-10). (D) Growth assay to test translation defects and its suppression by *RPB4/7* overexpression. The strains with the indicated genotypes were serially diluted in rich medium (YPDA) with or without 5 mg/mL neomycin.

Given that group 2 genes are involved in translation, ribosomal assembly, rRNA export and transport, this may indicate that phosphorylation of Rpb4 by Hrr25 could regulate the expression of genes whose products, including snoRNAs, are involved in processes such as translation. A direct role for Rpb4/7 in this process via mRNA imprinting has been also reported (Choder 2011; Dahan and Choder 2013; Duek et al. 2018; Chalabi Hagkarim and Grand 2020). For that reason, and because deficient Rpb4-P reduces Rpb4 dissociation from RNAPII at the 3′-end of genes, a prerequisite for the stalk binding to mRNA and export to the cytosol, we wondered if Rpb4 phosphorylation could be a mode to regulate mRNA imprinting. Thus, we analyzed the effect of the *rpb4-S/T-A* mutation in the subset of genes (∼1500 genes) whose mRNAs are imprinted by Rpb4/7, as shown by RNA immunoprecipitation sequencing (RIP-seq) (Garrido-Godino et al. 2020). Comparative metagene analysis of *wt* and the *rpb4-S/T-A* phospho-null mutant revealed a clear decrease in Rpb4 association with genes with imprinted mRNAs (Rpb4-RIP) compared with the rest of the genome (Fig. S7). This clearly shows that Rpb4 phosphorylation states can differentiate between genes encoding imprinted mRNA and the rest of the genes.

Altogether, our data show that Rpb4 phosphorylation has a role in transcription termination and that Rpb4 is a main substrate of Hrr25 kinase activity, which also modulates mRNA imprinting.

## DISCUSSION

Previous research has emphasized the importance of Rpb1-CTD phosphorylation in gene expression (Calvo and García 2012; Harlen and Churchman 2017); however, the potential role of phosphorylation in other RNAPII subunits has remained largely unexplored. In a recent proteomic study, five phospho-sites were identified in Rpb4: S125, S197, T134, T144, and T193 (Richard et al. 2021). To investigate their impact and to validate the biological significance of these phosphorylated residues, we generated two mutants, *rpb4-S/T-A* (phospho-null) and *rpb4-S/T-D* (phospho-mimetic). Although Rpb4-P accounts for approximately 5%-6% of the total Rpb4, mutating the five phospho-sites had a noticeable effect on cell growth, suggesting the relevance of Rpb4 phosphorylation. Specifically, the phospho-mimetic strain exhibited severe growth defects at 37°C, akin to cells lacking *RPB4* (*rpb4Δ*), probably due to a significant reduction in Rpb4-S/T-D protein levels (Fig. S1D). Furthermore, the Rpb4-Rpb7 interaction was impaired in *rpb4-S/T-D* cells, likely due to the negative charge of T144D that is located within at the Rpb4/7 interface (Fig. 1B and C). Rpb4-S/T-D/Rpb7 also exhibited assembly defect with RNAPII, probably because the integrity of the heterodimer is required for the interaction of both proteins with RNAPII (Sheffer et al. 1999). Mimicking Rpb4 hypo-phosphorylation also impaired cell proliferation. These seemingly paradoxical results support the idea that both phosphorylation states of Rpb4 are necessary for normal proliferation, suggesting that Rpb4 phosphorylation is a dynamic process and that phosphorylation is a transient modification. Interestingly, methylation of Rpb4 is also transient and dynamic (Richard et al. 2021). At present, it remains unclear whether the observed phenotypes in the *rpb4-S/T-D* mutant only results from the instability of the Rpb4/7 heterodimer (Fig. S1D) or from an additional mechanism that either promotes the dissociation of Rpb4/7 from RNAPII or hinders its assembly. This was not the case for the *rpb4-S/T-A* mutant, as Rpb4/7 stability remained unaffected and the integrity of RNAPII was not compromised. Hence, we conclude that defects in transcription or downstream processes following RNAPII assembly into genes, that characterize *rpb4-S/T-A,* are likely attributed to Rpb4 phosphorylation deficiency.

We have identified two enzymes that regulate the dynamic phosphorylation of Rpb4, Hrr25 and Fcp1. Both also act on the CTD: The kinase Hrr25 is known to phosphorylate CTD-Thr4 and plays a role in transcription elongation and termination (Nemec et al. 2019), and the phosphatase Fcp1 dephosphorylates the CTD (Ser2P and Thr4P) at the end of transcription to recycle RNAPII, but also has a role in transcription elongation (Cho and Buratowski 1999; Kobor et al. 1999; Mandal et al. 2002). Mutations in each enzyme resulted in similar phenotypes and exhibit genetic interaction with *rpb4-S/T-A* (Fig. 8D) or with *rpb4Δ* (Allepuz-Fuster et al. 2014). Moreover, overexpression of *RPB4/7* partially suppressed the growth defects of *hrr25-as*, indicating that Rpb4 is a major target of Hrr25 (Fig. 8D).

Rpb4 phosphorylation emerges as a novel mechanism for modulating transcription elongation and termination. As RNAPII progresses through the transcription cycle, the repertoire of PTM of the CTD changes (Buratowski 2003; Heidemann et al. 2013). Likewise, as demonstrated previously (Richard et al. 2021) and shown in the present study, the repertoire of Rpb4 changes as RNAPII proceeds through the transcription cycle. Rpb4 PTM are transient and specifically required for interactions with various factors involved in transcription (this paper), translation, or mRNA decay (Richard et al. 2021). Here we demonstrated that deficient Rpb4 phosphorylation led to transcription elongation defects (Figs. S2A and S3). Furthermore, Rpb4 phosphorylation was essential for proper transcription termination of snoRNAs (Fig. 7D) and for selecting the major polyadenylation site of *RNA14* (Fig. 7E). Thus, in cells lacking sufficient Rpb4 phosphorylation, promoter-proximal polyadenylation signals within *RNA14* were skipped less frequently, likely as a consequence of increased association of Rpb4 with the coding region (Fig. 3A and B), and a reduced elongation rate, which might facilitate the recognition of the weak signals by the cleavage and polyadenylation machinery (Geisberg et al. 2020). Specially, insufficient Rpb4 phosphorylation results in the highest increase of Rpb4 association with RNAPII at the 3′-ends. Therefore, Rpb4 phosphorylation might stimulate the dissociation of the stalk from RNAPII either before or during transcription termination. Interestingly, while writing this paper, it has been reported that the transcription factor Sfp1 accompanies RNAPII during transcription elongation and dissociates from RNAPII preferentially near the 3’ end of the transcription units (Kelbert et al. 2023). Because Rpb4 interacts physically and functionally with Sfp1, we speculate that both proteins dissociate from RNAPII near the 3′-end. On the other hand, in the phospho-null mutant, reduced association of Rpb3 with gene promoters (Fig. S3), as well as Rpb4 gene occupancy during initiation and early elongation (Figs. 3A and S3), are a consequence of stalk retention at the 3′-ends, and hence altered RNAPII recycling.

In summary, we propose that phosphorylation of Rpb4 regulates transcription elongation and facilitates both selection of the major polyadenylation sites and transcription termination of snoRNAs. We argue that Rpb4 phosphorylation stimulates dissociation of a portion of Rpb4/7 from RNAPII. This occurs mainly near the polyadenylation site, leading to mRNA polyadenylation and imprinting. Alternatively, in the case of genes whose transcription is regulated, for instance, by gene looping, Rpb4 needs to be dephosphorylated, similarly to Rpb1-CTD, to allow transcription reinitiation. Fcp1, which is required for proper dephosphorylation of the CTD after transcription termination and RNAPII recycling (Cho and Buratowski 1999; Kobor et al. 1999; Cho et al. 2001; Mandal et al. 2002; Allepuz-Fuster et al. 2014; Yurko and Manley 2018), may also dephosphorylate Rpb4 at 3′-ends to permit reassembly with RNAPII. Our results suggest that fine-tuning Rpb4-P states, regulated by Hrr25 and Fcp1, coordinate with CTD phosphorylation to control various transcription phases.

## MATERIALS AND METHODS

### Yeast strains and media

The strains used are listed in Table S1. Strain construction and other genetic manipulations were performed by following standard procedures (Burke 2000). The sequences of the oligonucleotides used in this study are available upon request.

### Generation of *rpb4* phospho-mutants

*RPB4* genes (*wt* and phospho-mutants) were obtained by gene synthesis and were introduced by homologous recombination into the *rpb4*::CaURA::TAP strain (Richard et al. 2021) to obtain the Rpb4-TAP strains (*wt*, *rpb4-S/T-A*, and *rpb4-S/T-D*) after selection on 5-Fluoroorotic acid containing media. The strains were then confirmed by PCR and Sanger sequencing.

### Analysis of Rpb4 phosphorylation *in vivo*

Cells were grown in 5 mL of rich medium to an optical density at 600 nm (OD_600_) of 0.5, harvested, washed, and then suspended in 20% trichloroacetic acid. To obtain whole cell lysates, cells were broken in a FastPrep homogenizer using glass beads. Proteins were then precipitated by centrifugation at 2200 *g* at 4°C for 5 min. The resulting pellet was solubilized in a solution containing 1 M Tris-HCl (pH 8) and Laemmli buffer. The sample was boiled for 10 min and centrifuged to remove insoluble material. The supernatant was loaded onto 10% SDS polyacrylamide gels containing Phos-tag (FUJIFILM Wako Chemicals Europe GmbH, Neuss, Germany) to facilitate the separation of Rpb4 phospho-bands. Then, proteins were transferred onto a 0.2 µm PVDF membrane, and western blotted with monoclonal anti-Rpb4 (clone 2Y14, Biolegend). In some cases, the Rpb3 level was determined as a loading control by using an anti-Rpb3 monoclonal antibody (clone 1Y26; Biolegend). ECL reagents were used for protein detection. The signals were acquired on film and with a ChemiDoc XRS system (Bio-Rad) and quantified with the Quantity One software (Bio-Rad), when required.

### Co-IP and western blot analysis

Two hundred milliliters of yeast cells were grown in rich medium to an OD_600_ of 0.8, harvested, washed with water, and lysis buffer (10 mM HEPES-KOH at pH 7.5, 140 mM NaC1, 1 mM EDTA, 10% glycerol, 0.1% NP-40, 1 mM PMSF, and 1x protease inhibitor cocktail [Complete; Roche]). The cell pellet was flash frozen in liquid nitrogen with 1.5 mL of lysis buffer and grounded in a Spex Freezer Mill 6775 to a fine powder. Afterwards, the cell lysate was thawed on ice and centrifuged at 13200 rpm for 20 min. The supernatant was collected and the total protein concentration was estimated. To immunoprecipitate RNAPII core or Rpb4, 1 µg/µL of anti-Rpb3 or anti-Rpb4 antibody was incubated with 25 µL of magnetic beads (Dynabeads M-280 Sheep anti-Mouse IgG, 11202D, Invitrogen) for 1 h at 4°C in phosphate-buffered saline (PBS) with 0.1% bovine serum albumin (BSA). After washing, the antibody coupled to beads was incubated with 5 mg of protein for 2 h at 4°C. Rpb4-TAP was immunoprecipitated by incubating 5 mg of proteins with IgG beads (Dynabeads Pan Mouse IgG) for 3 h at 4°C. All the immunoprecipitated samples were extensively washed with lysis buffer and beads were suspended in SDS-PAGE sample buffer. Thereafter, they were incubated at 65°C for 10 min and then the supernatants were loaded onto an SDS-PAGE gel.

Western blot analysis was performed using the appropriate antibodies in each case, acquired from the following vendors: anti-phosphoglycerate kinase (Pgk1, 459250; Invitrogen), anti-Rpb1 (8WG16, Covance), anti-Rpb3 and Rpb4 (Biolegend), anti-CTD-Ser2P (ab5095, Abcam), anti-CTD-Ser5P (clone 4H8, Millipore), and anti-Thr4P (clone 6D7, Active Motif). ECL reagents were used for detection.

### ChIP-qPCR and ChIP-seq

ChIP with the corresponding antibodies and DNA purification followed by qPCR were performed as described previously (Allepuz-Fuster et al. 2014; Garavis et al. 2017), with minor modifications. The chromatin was sheared in a Bioruptor Plus (Diagenode) and then immunoprecipitated using Dynabeads coupled to the corresponding antibody. qPCR was performed in triplicate using the DNA of at least three independent ChIP experiments in a CFX96 Detection System (Bio-Rad). Quantitative analysis was carried out with the CFX96 Manager Software (version 3.1, Bio-Rad). The values obtained for the immunoprecipitated PCR products were compared with those of the total input, and the ratio of the values from each PCR product from transcribed genes to a non-transcribed region of chromosome XII was calculated. In some cases, ChIP experiments were performed in parallel either with no antibodies or, in the specific case of Rpb4 ChIP, using chromatin from the *rpb4Δ* strain as a negative control. The numbers on the y-axis of the graphs are detailed in the corresponding figure legends.

For ChIP-seq experiments, purified DNA was also submitted for deep sequencing. Each experiment was carried out in three biological replicates. Library DNA was prepared from immunoprecipitated DNA and its corresponding input DNA following the manufacturer’s instructions, and was sequenced in an Illumina HiSeq 2500 system to an average output of 15 (1 × 50 nucleotides [nt]) million reads. Sequencing fastq files were inspected for general quality issues with *FastQC* (http://www.bioinformatics.babraham.ac.uk/projects/fastqc) and *MultiQC* (Ewels et al. 2016). *Trimmomatic* (Bolger et al. 2014) was used for the initial clipping of standard Illumina adapters. Then, low-quality reads were removed in a two-step manner: The sliding window trimming function was used to remove parts of reads with qualities below 20 (using a window width of 4 bases), and the remaining reads that were < 20 nt were dropped. The remaining high-quality reads were aligned to the *sacCer3* reference genome with *Bowtie2* (Langmead and Salzberg 2012), using default parameters. Average metagene analysis of ChIP-seq datasets was performed with the R package *ngs.plot* (Shen et al. 2014). To visualize only one profile per sample, the biological replicate alignment files of each sample were merged with the function merge from *samtools* (Danecek et al. 2021), and then the merged files were re-indexed. The metagene plots from *ngs.plot* were drawn using the following parameters: *-G sacCer3 -FL 300 -L 500 -RB 0.05 -SE 0*. Heatmaps of the Rpb4/Rpb3 signal were also drawn with *ngs.plot*, using the additional parameter *-SC global* to obtain the same color scale for all plots. Density scatterplots were drawn with the *ggpointdensity* package (https://GitHub.com/LKremer/ggpointdensity) in *ggplot2*, and the Spearman correlation values were added to the plots with the *ggpubr* package (https://rpkgs.datanovia.com/ggpubr).

### RNA purification and RT-PCR

Total RNA was extracted as described in a previous study (Schmitt et al. 1990) or by using the Qiagen RNeasy RNA isolation kit for samples submitted to deep sequencing. RT-PCR was performed using the iScript cDNA Reverse Transcription Supermix Kit (Bio-Rad), following the manufacturer’s instructions. RT-PCR was performed in triplicate with at least three independent cDNA samples.

### RNA-seq of *wt* and phospho-mutant strains

Total RNA was extracted from the corresponding strains from three biological replicates and subjected to rRNA depletion. The rRNA-depleted fraction was used to construct libraries that were sequenced in an Illumina HiSeq 2500 system to an average output of 12 (1 × 50 nt) million strand-specific reads. The pre-processing and removal of low-quality reads from RNA-seq files was identical to that of ChIP-seq files described above. High-quality reads were aligned to the *sacCer3* reference genome with *HISAT2* (Kim et al. 2019), using default parameters. Raw read counts were extracted from alignment files with *featureCounts* (Liao et al. 2014), using the *R64-1-1.38.ggf3* annotation file from Ensembl. DE analysis was carried out with the *Limma-Voom* method (Law et al. 2014), using the raw reads extracted with *featureCounts* from each biological replicate and filtering genes with < 1 read per million total mapped reads in each individual sample. Genes were considered to be DE if they had an adjusted *p*-value (Benjamini-Hochberg method) of < 0.05. Comparative GO enrichment analysis was performed with the *compareCluster* function from the R package *clusterProfiler* (Wu et al. 2021), using the *org.Sc.sgd.db* annotation and the biological process ontology. GO term redundancy was reduced with the *simplify* function that was implemented with the following parameters: *cutoff* = 0.5, *by* = “p.adjust,” *select_fun =* min, *measure* = “Wang,” *semData* = NULL. Once the *compareCluster* object was created, the results were visualized with the *dotplot* function from the R package *enrichplot* (https://yulab-smu.top/biomedical-knowledge-mining-book/). Average metagene analysis of snRNAs was generated by using *ngs.plot* with the following parameters: *-G sacCer3 -FL 50 -L 250 -RB 0.05 -F misc -SE 0*.

Reanalysis of the *hrr25-is* mutant RNA-seq datasets from (Nemec et al. 2019). Two *wt* and two *hrr25-is* mutant raw sequencing files were retrieved from the ENA repository (stored under accession code PRJNA419200). Replicates were pre-processed with *Trimmomatic* to remove adapters and low-quality reads and then aligned to the *sacCer3* reference yeast genome with *TopHat2*, using default parameters. Raw counts were obtained using *featureCounts*, and DE analysis between *mutant* and *wt* samples was performed with *DESeq2* (Love et al. 2014), using default parameters. Genes were considered to be DE if they had adjusted *p*-values (false discovery rate [FDR] method) of < 0.05.

### Purification of rRpb4/Rpb7-6His

The Rpb4/7 complex was purified from *Escherichia coli* expressing a plasmid containing Rpb4 and Rpb7 with a six-histidine tail fused to its C-terminus, as described in (Garavis et al. 2017). rGST-CTD was expressed and purified as described previously (Garcia et al. 2010).

### Pull-down assays

Recombinant Rpb4/Rpb7-6His protein (5 μg) was incubated in a 50 μL slurry of HisPur Cobalt Resin (Thermo Scientific) in binding buffer (20 mM Tris [pH 7.5] and 50 mM NaCl) for 1 h at 10°C. Thereafter, the resin with attached Rpb4/Rpb7-6His was washed three times with binding buffer and then WCE from a *rpb4Δ* strain was added to the resin and incubated for 2 h at 10°C in binding buffer. After incubation, the resin was washed four times with washing buffer (20 mM Tris [pH 7.5] and 50 mM NaCl) and then proteins were eluted with elution buffer (20 mM Tris pH 7.5, 50 mM NaCl, and 300 mM imidazole) for 10 min at room temperature. A fraction of the eluate (10%) was analyzed by SDS-PAGE and the rest (90%) was precipitated with trichloroacetic acid and submitted to MS analyses. The pull-down assay was performed in triplicate with three independent WCE.

The detection and identification of proteins was carried out as follows. The samples were digested with trypsin, using S-Trap columns. The tryptic peptides were cleaned with C18 ZipTip and the equivalent of 750 ng (quantified by QBIT) was analyzed by liquid nanochromatography coupled to MS, in DDA mode, using a Thermo Ultimate 3000 liquid chromatograph and a Thermo Orbitrap Exploris mass spectrometer OE240. The flow of the liquid chromatograph was 250 nL/min; the C18 reverse-phase column used had an internal diameter of 75 µm and a length of 25 cm; and the total length of the gradient was 60 min. The obtained MS1 and MS2 spectra were used to launch a search, using the Mascot search engine, against a combined database consisting of 6118 *S. cerevisiae* entries. The parameters used for the search were: a tolerance of 10 ppm and 0.02 Da for the precursor ions and fragments, respectively; carbamidomethylation in cysteines as a fixed modification; acetyl at the N-terminus of the protein and methionine oxidation, Gln->pyro-Glu (N-term Q), and Glu->pyro-Glu (N-term E) as modification variables. Finally, the identified proteins were quantified by using the exponentially modified protein abundance index (EmPAI) as described previously (Ishihama et al. 2005).

### *In vivo* inhibition of *hrr25-as* and *hrr25-is* kinase activity

A 200 mL culture of cells expressing Hrr25-TAP (*wt* or *hrr25-as*) was grown in YPD media at 28°C to an OD_600nm_ of 0.3. The culture was split into two halves and treated with 3MB-PPI inhibitor (Toronto Research Chemicals) at a final concentration of 20 μM (or an equivalent volume of DMSO) for another 1 h. Then, cells were collected, washed, and flash frozen in liquid nitrogen for further WCE preparation in a freezer mill (see below). For the assays performed with the *hrr25-is* strain, cells were grown similarly, but 20 μM CMK (MedChem Express) was used instead of 3MB-PPI.

### Purification of TAP-tagged kinases

Strains expressing TAP-tagged kinases (Kin28, Bur1, and Hrr25) were grown in 200 mL of rich medium to an OD_600_ of 1.0, harvested, washed with water, and then suspended in 500 μL of tandem affinity purification buffer A (20 mM HEPES pH 7.9, 300 mM potassium acetate, 0.5 mM EDTA [pH 8.0], 10% glycerol, and 0.05% NP40) containing 1 mM DTT and 1 mM protease and phosphatase inhibitors. The cell suspension was flash frozen in liquid nitrogen and then ground in a Spex Freezer Mill 6775 to a fine powder. Afterwards, the cell lysate was thawed on ice, centrifuged, and the supernatant recovered. Fifty microliters of a 50/50 slurry of IgG-Sepharose 6 Fast Flow beads (GE Healthcare) in tandem affinity purification buffer A was added to the supernatant. After incubating for 3 h at 4°C, the beads were pelleted and washed five times with tandem affinity purification buffer A (500 μL each). The corresponding kinases were eluted overnight at 4°C in 25 μL of tandem affinity purification buffer A with 1 mM DTT and 10 U of AcTEV (Invitrogen). The purified kinases were pooled and concentrated using Millipore centrifugal devices in buffer D kinase (20 mM HEPES [pH 7.5], 10% glycerol, 2.5 mM EGTA, 15 mM magnesium acetate, and 100 mM potassium acetate).

### *In vitro* kinase assays

Each assay was performed with 5 µg of tandem affinity purified kinase and 300 ng of rRpb4/7×6His (or 100 ng of GST-CTD) in a 30 µL of buffer D kinase (20 mM HEPES [pH 7.5], 10% glycerol, 2.5 mM EGTA, 15 mM magnesium acetate, and 100 mM potassium acetate) containing protease and phosphatase inhibitors, 1 mM DTT, and 0.5 µL γ-ATP (250 µCi), and 0.5 µL of 50 mM cold ATP for the radioactive assay or 1 mM cold ATP for the nonradioactive assays. Reactions were performed at 30°C for 2 h and stopped with Laemmli loading buffer. Thereafter, they were incubated at 65°C for 5 min and loaded onto an SDS-PAGE gel, and then transferred to a nylon membrane. Radioactive signal was acquired using a PhosphorImager and then incubated with either anti-Rpb4 or anti-Rpb1 and anti-Ser5P antibodies.

## Statistical analysis

All tests were performed in triplicate (n = 3). Prior to analysis, the data were normalized with the square root and, later, scaled by the Pareto method to avoid differences introduced by the units of measurement (Seisonen et al. 2016). Data processing was performed with the statistical package Statgraphics Centurion XVI.II (Scientific Time Sharing Corporation, Rockville, MD, USA). The applied methodology was one-way analysis of variance (ANOVA), and Fisher’s test was used to establish homogeneous groups at a significance level of *p* ≤ 0.05.

## Data availability

All data needed to evaluate the conclusions in this study are present in here and/or the in the Supplemental Material.

Processed data were deposited to GEO (GSE243009).

## Acknowledgments

We are grateful to S. Camero, Dr. J.M. Pérez-Cañadilla, and Dr. J.M. Pereda for assistance in rRpb4/7 expression and purification; Dr. Ansari for *hrr25* strains and Dr. Boone for *fcp1-5001* mutant. We also thank the Proteomic Department of the CNB-CSIC, the Genomic Facility at the CRG, and Dr. J.E. Pérez-Ortín for helpful comments on the manuscript. The O.C. laboratory is supported by the grant PID2020-116396GB-I00 funded by MCIN/AEI/10.13039/501100011033.

## Author contributions

O.C. designed all the experiments, and initiated and led the project. A.G-J., C.G-J., I.M, MJ.A., and O.C. performed the experiments. A.G-J., C.G-J., I.M., and O.C. analyzed most of the data. A.J. analyzed all deep sequencing data and made the corresponding figures. C.F-T. made figures for structures and helped interpret the protein interaction data. S.R. and M.C. provided valuable information on yeast strains and plasmids to initiate the project. O.C. wrote the manuscript with the input from all the authors, especially M.C.

